# Neural networks with optimized single-neuron adaptation uncover biologically plausible regularization

**DOI:** 10.1101/2022.04.29.489963

**Authors:** Victor Geadah, Stefan Horoi, Giancarlo Kerg, Guy Wolf, Guillaume Lajoie

**Affiliations:** Program in Applied and Computational Mathematics, Princeton University, Princeton, U.S.A.; Mila - Quebec Artificial Intelligence Institute, Montréal, Canada; Département de Mathématiques et Statistiques, Université de Montréal, Montréal, Canada; Département d’Informatique et Recherche Opérationelle, Université de Montréal, Montréal, Canada; Canada CIFAR AI Chair

## Abstract

Neurons in the brain have rich and adaptive input-output properties. Features such as heterogeneous f-I curves and spike frequency adaptation are known to place single neurons in optimal coding regimes when facing changing stimuli. Yet, it is still unclear how brain circuits exploit single-neuron flexibility, and how network-level requirements may have shaped such cellular function. To answer this question, a multi-scaled approach is needed where the computations of single neurons and neural circuits must be considered as a complete system. In this work, we use artificial neural networks to systematically investigate single-neuron input-output adaptive mechanisms, optimized in an end-to-end fashion. Throughout the optimization process, each neuron has the liberty to modify its nonlinear activation function, parametrized to mimic f-I curves of biological neurons, and to learn adaptation strategies to modify activation functions in real-time during a task. We find that such networks show much-improved robustness to noise and changes in input statistics. Importantly, we find that this procedure recovers precise coding strategies found in biological neurons, such as gain scaling and fractional order differentiation/integration. Using tools from dynamical systems theory, we analyze the role of these emergent single-neuron properties and argue that neural diversity and adaptation play an active regularization role, enabling neural circuits to optimally propagate information across time.

## 1 Introduction

Biological neurons have diverse input responses and adaptation properties (Gjorgjieva et al., 2016; Weber et al., 2019). How the rich dynamics of biological neurons combine with network interactions to support complex tasks, such as sensory integration and behavior, remains largely unresolved. While the past decades have seen considerable work aimed at elucidating single neuron coding properties, most efforts have been “bottom up”, modeling mechanistic features observed in biology and analyzing their computational impact. We argue that to shed light on the system-level role of single neuron properties, a “top-down” approach is needed. One way to achieve this is with deep-learning optimization, where “goal-driven” models aim to solve system-level objectives, and emergent neuron properties are studied. In recent years, this method has been extremely successful in capturing single neuron static tuning properties, such as that in the visual system (Yamins and DiCarlo, 2016). In this work, we use a goal-driven approach to investigate adaptive input-output properties of neurons that emerge from end-to-end optimization of recurrent neural networks (RNNs), and shed light on their role in biological systems.

A central dynamic component of single-neuron coding is the transformation of input currents into output firing rates neuron execute, as measured by so called *f-I curves*, or *activation functions* (AF). These are both adaptive and diverse across neurons. At the heart of this modularity lies the efficient coding hypothesis, a theoretical paradigm by which neurons aim to be maximally informative about the inputs they encode (Barlow, 1961; Laughlin, 1981). Supported by this principle, neurons are known to effectively modulate their f-I curve in response to constant step-like stimulus, in a process know as *spike frequency adaptation* (SFA) (Benda and Herz, 2003). It has been shown that SFA and other adaptive mechanisms in single neurons enable faithful encoding of input signals regardless of stimulus baseline, a crucial feature for animals subject to changing environments (Fairhall et al., 2001; Peron and Gabbiani, 2009; Gjorgjieva et al., 2016). SFA also facilitates information integration over long timescales (Pozzorini et al., 2015), and provides robustness to rapid variation and noise (Lundstrom et al., 2008). At the network level, adaptive neural responses have been shown to support efficient coding with metabolic advantages (Gutierrez and Denève, 2019), facilitate computations over long timescales (Bellec et al., 2018; Fitz et al., 2020; Salaj et al., 2021), and even enable forms of Bayesian inference (Deneve, 2008; Kilpatrick and Ermentrout, 2011). Recent work also shows robustness gains from learned modulated neural dynamics (Vecoven et al., 2020) and with diverse and dynamics synapses and f-I curves (Burnham et al., 2021; Winston et al., 2022), While a number of coding advantages of diverse and dynamic single neuron responses are now established, it is still unknown how these mechanisms have come to bear, and how they influence learning and configuration of larger neural networks that support system-level tasks such as perception or prediction.

In parallel, modern artificial neural networks used in artificial intelligence (AI) loosely mimic neural responses with simple AFs (also called *nonlinearities*) which transform summed inputs to an artificial neuron into a scalar state value, akin to a firing rate. While different shapes of activation functions have been used, and even optimized (Hayou et al., 2019), the prevailing sentiment in AI is that a simple AF such as the *rectified linear unit* (ReLU) is enough for large networks to implement almost any transformation. In fact, this is mathematically guaranteed by the universal function approximation theorem, stating that large enough nonlinear neural networks can implement any function (Cybenko, 1989; Hornik et al., 1989). Reconciling the diverse and dynamic nature of biological neurons’ input-output properties with the computational function of the large networks in the mammalian brain, for example, is therefore a tricky exercise. The prevalent hypothesis is that the single neuron input-output richness found in the brain has evolved and been optimized to guide network-level function such as stable population dynamics, and coordinated learning.

In this work, we propose a step towards complementing longstanding mechanistic investigations into f-I-response diversity and adaptation, through the lens of goal-driven optimization. Using simple artificial neural networks and deep learning, we ask: given the possibility to implement a wide range of single neuron input-output properties, including rapid adaptive mechanisms, do networks optimized end-to-end develop biologically realistic solutions at the single neuron level? If so, can we reconcile single-neuron properties with network-level mechanisms? To address this, we concentrate on the problem of perception on sequential stimuli, such as visual input streams. Our goal is to prescribe the simplest recurrent neural network (RNN) possible that has enough flexibility to develop optimal solutions for its units’ AFs. As such, we propose a two-parameter family of AFs mimicking the diversity of f-I curves that can be implemented by known neural types, and interpolating between often used nonlinearities in AI. In addition, we implement a dynamic controller that modulates AFs in real time, acting locally and independently at each neuron. This controller, implemented with a distinct and smaller RNN, models the genetically encoded adaptation strategy that would have been refined by evolution (see e.g. Sandler et al. (2021); Vecoven et al. (2020) for similar ideas). We then train this system end-to-end on sequential classification tasks. We call our novel adaptive artificial neuron *Adaptive Recurrent Unit* (ARU).

Our findings can be summarized in three points. First, we find that both diverse and adaptive AFs help the main RNN learn tasks, and provide surprising robustness to noise and distractors. Second, we investigate the learned solutions obtained by the optimization procedure and find that surprisingly, a number of biologically realistic strategies are implemented. Indeed, optimal AFs take on biologically plausible configurations (i.e. not simple sigmoid or ReLU), diversity of AFs is an important and necessary feature for robustness, and crucially, the adaption controller implements gain scaling and fractional order integration and differentiation, just like several neocortical neurons. Finally, we analyze the optimization mechanism that led to these solutions and find that diversity and adaptation acts as a dynamic regularizer, enabling the main RNN to remain in a regime close to the edge of chaos where information transmission and error gradients propagate optimally.

## 2 Results

### 2.1 Flexible activation functions in recurrent neural network models

In line with the goal of isolating the role of activation functions, we elect to use a simple “vanilla” RNN model for experiments. The vector equation for the recurrent unit activation 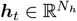 in response to input 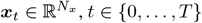 is given by

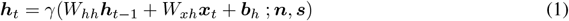

where the output 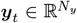 is generated by a linear readout ***y***_*t*_ = *W*_*hy*_***h***_*t*_ + ***b***_*y*_. Weight matrices *W*_(·)_ and biases *b*_(·)_ are optimized via gradient descent and backpropagation through time (BPTT) in all experiments (see Methods). Departing from standard RNNs, the AF (or nonlinearity) *γ* comes from a differentiable family parametrized by

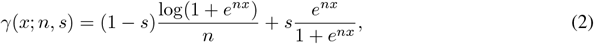

for scalar input *x*, with two parameters controlling its shape: the **degree of saturation** *s* and **neuronal gain** *n*^1^. This is a *s*-modulated convex sum of two differentiable functions: the non-saturating softplus (*s* = 0), and the saturating sigmoid (*s* = 1), while *n* rescales the domain and controls response sharpness, or gain (Sompolinsky et al., 1988). Fig. 1**a** shows the graph of *γ* for different values of (*n, s*), interpolating between well-known activation functions in deep learning. This also captures key properties of f-I curve shapes present in different neuronal types. For instance, Type I neurons show gradual frequency increase with increasing applied current (i.e. *γ* with low *n, s* ≥ 0), whereas Type II neurons show sharp firing onset at non-zero frequencies (i.e. *γ* with high *n*, and low *s >* 0). We let each neuron have their private AF, with activation parameters {*n*^*i*^, *s*^*i*^} associated with neuron *i* (see Methods §B.1 for a comparison to when AF parameters are shared). As *γ* is differentiable in both *s* and *n*, one can include these parameters in the optimization scheme. We consider two main learning frameworks for analysis: (1) *static*, and (2) *adaptive* AFs (see Figure 1**b** for a schematic). **(1) Static**: each neuron’s AF is optimized by gradient descent (see Methods §5.3) but remains static throughout neural dynamics during input processing (Fig. 1**c**). **(2) Adaptive**: the AF of each neuron *i* has parameters 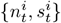 varying in time *t*, governed by parameterized mechanisms optimized through gradient descent.

**Figure 1.**
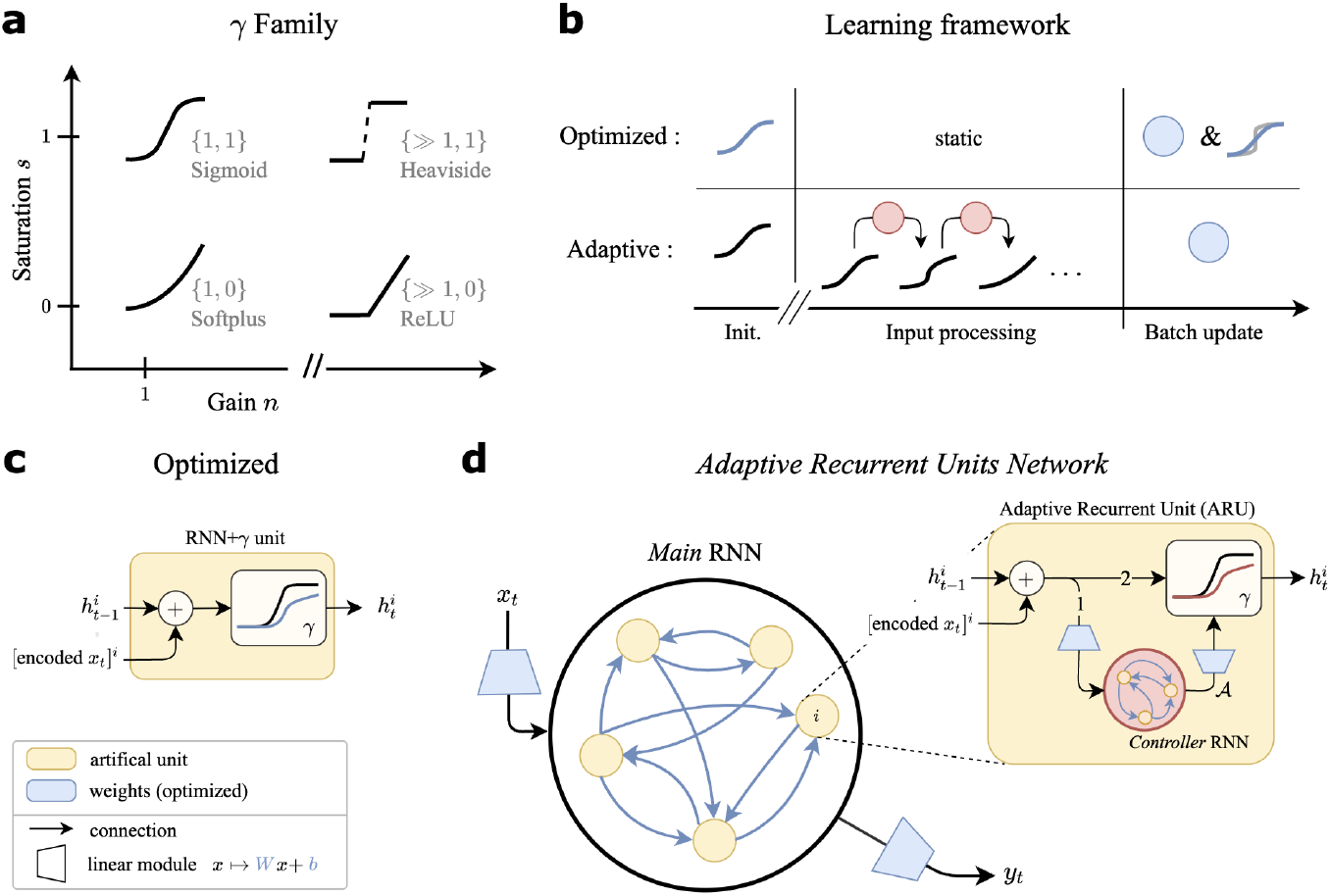
Model details. (**a**) Various shapes of the *γ* activation function (AF), represented in the parameter space {*n, s*}. We attain commonly used nonlinearities. (**b**) Different settings considered for modulation of the AF. This modulation is either done alongside training (*Optimized*) or online (*Adaptive*). Blue indicates learned parameters; notice the blue activation function in the *Optimized* setting, updated alongside weights. (**c**) Artificial unit *i* of a standard RNN with *γ* activation function. Legend at the bottom applies to (**c**-**d**). (**d**) Graphical depiction of the Adaptive Recurrent Units (ARU) and associated recurrent network model. Numbers {1, 2} on the arrows in the ARU represent the order of processing. Removing the adaptation mechanism **𝒜**, we recover the RNN+*γ* model in **c**. Each neuron has a private copy of the controller RNN (in red).

In the adaptive setting, our goal is allow for **adaptation mechanisms** to be discovered by optimization, mapping neural pre-activations 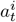 to a nonlinear activation function 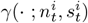, akin to spike-frequency adaptation in cortical neurons. To this end, we train an **adaptation controller** to produce the dynamics of 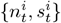 (Fig. 1**d**). This controller is itself an RNN module, chosen because of its universal function approximation properties, and does not represents neural circuits like the main RNN. The adaptation controller takes in the presynaptic inputs to a neuron *i*, and returns AF parameters via linear readouts at each time step. Crucially, each neuron has an identical copy of this controller RNN (shared parameters), but each copy controls its single neuron independently of the others. The equations for the controller RNN are given by

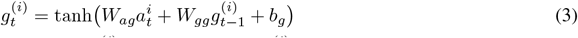

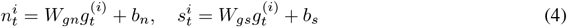

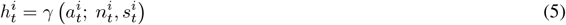

where 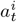 is the presynaptic input to neuron *i* at time step *t* (given by 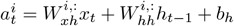 following Eq. 2), and 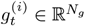 is the vector of hidden states for the controller network associated to neuron *i* at time *t*. The controller parameters Θ_**𝒜**_ = {*W*_*ag*_, *W*_*gg*_, *W*_*gc*_, *b*_*g*_, *b*_*c*_} are shared across all neurons *i* = 1, …, *N*_*h*_, and are optimized end-to-end (see Methods for details). The adaptation controller is thus a shared mechanism across neurons, but with each neuron having a private copy for local control. This is similar to the shared genetic information governing complex ionic machinery, but which individual neurons operate independently from others.

### 2.2 Neural adaptation and diversity improves RNN performance and robustness to input perturbations

We use basic perceptual classification tasks to explore static and adaptive AF optimization in our RNN: (1) sequential classification: MNIST (Le et al., 2015) digits from a permuted sequential sequence of pixels (**psMNIST**), and (2) a grayscaled and sequential version of the CIFAR10 classification task (**sCIFAR10**). In both cases, an image is presented to the RNN one pixel at a time, and class membership is inferred at the end (digit number for sMNIST, and “truck”, “plane”, “horse”, etc. for sCIFAR10). See Methods for further details on both tasks. As baselines models for comparison, we use a standard RNN+ReLU (Glorot et al., 2011), as well as the gated LSTM (Hochreiter and Schmidhuber, 1997) and GRU (Cho et al., 2014) architectures. The latter two models are known to be more efficient at learning long time-dependencies thanks to architectures that leverage gates. This is, however, not biologically plausible since it relies on non-local mechanisms. Nevertheless, these models offer a good comparison to highlight the advantages of optimized single neurons AF mechanisms which are biologically plausible. As the sCIFAR10 task is more difficult, we also consider a scenario where a convolutional encoder is trained to pre-process inputs. This is optimized end-to-end for each model compared. In all model comparison, we aim to maintain comparable parameter counts (see Methods)

#### Optimized activation functions improve performance

First, we assess model performance by their classification accuracy on held-out test data. We find that the introduction of optimized *γ*(·;; *n, s*) AF, both static and adaptive, provides a considerable increase in performance compared to baselines (see Table 1). On the psMNIST task, RNN+*γ* (static) outperform both ReLU and gated (LSTM, GRU) baselines. In this testing setting, we did not observe a significant difference between the RNN+*γ* and the adaptive ARUN. Similar results are obtained on the sCIFAR10 task. First using only a linear encoder module like the psMNIST task, we observe that while the GRU offers the highest performance by far, the optimized RNN+*γ* achieves greater classification accuracy and provides a significant improvement over the RNN+ReLU and LSTM. The introduction of a convolution encoder module significantly increased all the networks’ performance. With this encoding scheme for the sCIFAR10 task, all models offer similar performance.

**Table 1:** Test accuracy on held-out data on the permuted sequential MNIST and sequential CIFAR10 classification tasks. Performance is evaluated on held-out standard input samples (“Original”), and on those same inputs but with a noisy step perturbation (***ξ***_*t*_ 𝒩(**5**, 2^2^*I*) for psMNIST, ***ξ***_*t*_ 𝒩(**1**, *I*) for sCIFAR10) applied to the neurons directly (“Noise Pert.”, see Fig. 2**a**-top). Mean and standard deviation over three random seeds.

| (%) | psMNIST |  | sCIFAR10 |  |  |
| --- | --- | --- | --- | --- | --- |
|  | Original | Noise Pert. | Original |  | Noise Pert. |
| Model |  |  | Lin. encoder | Conv. encoder |  |
| RNN+ReLU | 90.1 ± 0.2 | 33.0 ± 18.9 | 28.1 ± 0.2 | 75.1 ± 2.0 | 50.0 ± 9.9 |
| LSTM | 89.6 ± 2.6 | – | 36.4 ± 2.3 | 73.9 ± 1.5 | – |
| GRU | 93.3 ± 0.2 | – | 61.3 ± 0.4 | 74.2 ± 2.0 | – |
| RNN+$\gamma$ (static) | 95.4 ± 0.5 | 84.0 ± 9.1 | 42.5 ± 3.1 | 75.4 ± 1.9 | 54.2 ± 2.4 |
| ARUN | 95.4 ± 0.2 | 94.7 ± 0.4 | 44.3 ± 2.9 | 75.1 ± 2.2 | 52.3 ± 9.6 |

#### Adaptive units improve robustness to changes in environment

We now study model robustness to perturbations unseen during training. First, we draw inspiration from optogenetic stimulation and inject an external drive *ξ >* 0 directly to each neuron before the AF is applied during *τ* time-steps (Fig. 2**a**-top). Such external drive can either be noisy, drawing 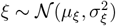 i.i.d. across neurons and in time from normally distributed noise of non-zero mean *μ*_*ξ*_ *>* 0 and *σ*_*ξ*_ such that *ξ* is almost surely positive, or non-random at a fixed amplitude *ξ >* 0. For those step perturbations we do not consider the LSTM and GRU architectures, as there is no parallel to injecting noise to the main network due to their multiple gating features. Second, we transform the network-level inputs *x*_*t*_ by applying a sinusoidal change in contrast (Fig. 2**a**-bottom), thus altering the input statistics directly. More details in Methods §5.5.

**Figure 2.**
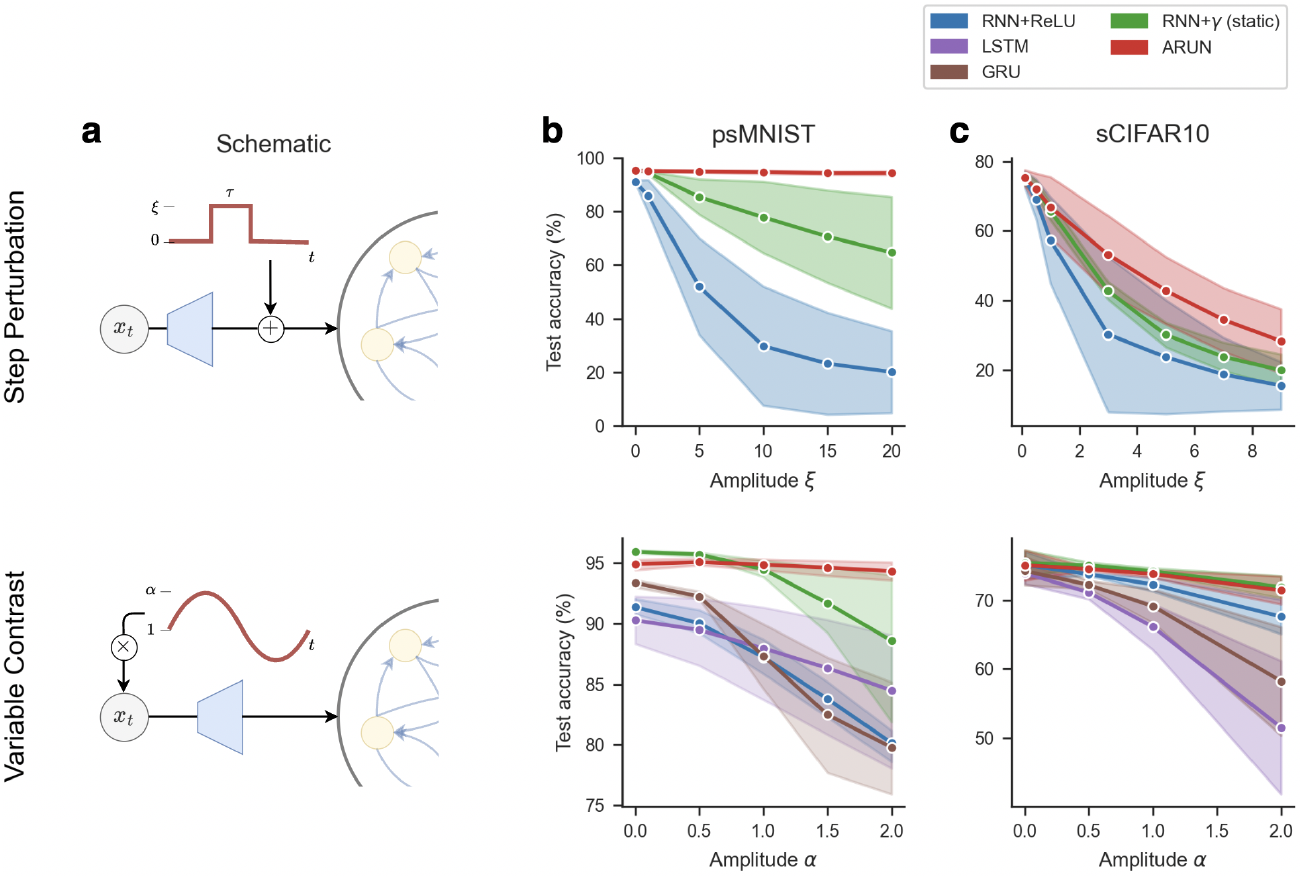
Performance and robustness of RNN architectures on sequential classification tasks. (**a**) Schematic of the perturbations considered, either an additive step perturbation of amplitude *ξ >* 0 for *τ* time-steps (top), or a multiplicative change in contrast applied to the pixels inputs (bottom). Performance as test accuracy on held out data on the permuted sequential MNIST (**b**) and sequential CIFAR10 (**c**) classification tasks. For the additive step perturbation analysis (top), we fixed the step perturbation period at *τ* = 200 time-steps starting at *t* = 200, and plot the performance as a function of the drive amplitude *ξ*. Similar qualitative results hold for noisy step perturbations (see Table 1). For the multiplicative variable contrast analysis (bottom), we consider a sinusoidal transform of fixed phase 0, of frequency of 1.0 for psMNIST and 2.0 for sCIFAR10 (best overall performance accross networks) and plot the performance as a function of the transform amplitude *α*.

We observe that networks with adaptive nonlinearities provide an increased ability to mitigate the changes in input statistics from the transformed digits or in response to an added stimulus. In psMNIST, the ARUN outperforms other architectures on the noisy drive (Table 1), and shows high robustness to variable drive amplitude (Fig. 2**b**-top). For varying amplitude on the multiplicative contrast experiment, we observe that the RNN+*γ* model already offers improved robustness, second to again the ARUN showing the lowest decrease in performance. Performance experiments on the sCIFAR10 task slightly less conclusive, as expected, attributable in part to the overall lower performance for all networks. We nonetheless observe that the ARUN still offers more robustness to step perturbations (Fig. 2**c**-top), and overall lessend decrease in performance on the variable contrast (Fig. 2**c**-top) with respect to baselines.

On top of the results presented in Figure 2, we conducted a sensitivity analysis with respect to the various parameters of the transformations (see Appendix Figures 13-16). First, we varied the phase and amplitude of the sinusoidal transformation applied on inputs, and we observe that the ARUN presents the best robustness on the psMNIST task. Second, we varied the amplitude and length of the step-drive applied on neurons. In this driven case, again on the psMNIST task, the ARUN presents a test loss of an order of magnitude lower than the other RNN models while varying the parameters. Similar sensitivity analysis on sCIFAR10 suffers from a similar lack of significant differences between models as the results presented previously. We still observe similar trends towards the beneficial impact of our adaptive architecture. Thus in all, endowing networks with adaptive nonlinearities presents advantages in mitigating changes in input statistics.

### 2.3 Top-down optimization of adaptive RNNs recovers biological dynamic coding mechanisms of single neurons

When trained on temporal perception tasks (sequential MNIST/CIFAR10), we demonstrated that our network of ARUs shows improved robustness to noise and changes in input statistics. In this section, we investigate the adaptive strategies learned at the single neuron level, and draw parallel with those found in the brain. We find that ARU neurons recover three key ingredients found in biological neurons: (1) heterogeneity, (2) gain scaling, and (3) fractional differentiation/integration. We endowed our RNNs with adaptive capabilities through minimial inductives biases, and they could have, in principle, chosen any adaptive of AF strategy, biologically realistic or not.

#### Heterogeneity

Heterogeneous activation functions already provide a setting more reminiscent of the diversity of activations in cortical networks. In Section §2.1 we demonstrated that heterogeneity was beneficial to task performance and robustness (albeit not significantly, see Appendix §B.1). Furthermore, we observe that when the activation function *γ* is initialized homogeneously, the optimization procedure leads to heterogeneity in the activation functions across the network (Fig.3**a**-top). Ref. (Winston et al., 2022) show similar results when AFs are parametrized following known relations between ionic currents and f-I curves. Further experiments (details included in Appendix §B.3) consider trained RNN+*γ* networks reading psMNIST digits rotated by 45*°*. As opposed to the perturbation experiments highlighted previously, this also changes the temporal order in which the inputs are fed. In this new setting, we observed an increase in {*n, s*} heterogeneity upon changes in task temporal statistics (Fig.3**a**-bottom), when all other parameters are kept fixed. Simply allowing the AFs to be modulated could recover over a quarter of lost performance (over 25%) in this altered task. These results show that heterogeneity is both beneficial and is learned through optimization.

#### Adaptation implements gain scaling

As a first indication of ARUs implementing optimal coding mechanisms akin to biological neurons, we find the general gain adaptation behavior to follow general gain scaling principles of cortical neurons (Laughlin, 1981; Fairhall et al., 2001). We subjected ARUs to a low-noise signal 𝒩 (0, 0.01^2^) for *t* = 200 time steps, followed by a i.i.d samples from 𝒩 (0, *ξ*^2^) for varying *ξ >* 0 during another *t* = 200 time steps, before returning to the original low noise (see inlet Fig. 3**b**). We observe the steady-state gain *n* (*ξ*) to display a power-law dependence on *ξ* (Fig. 3**b**-left) (Lundstrom et al., 2008). As for the saturation, we observed a exponential dependence on *ξ* (Fig. 3**b**-right). These have the combined effect of following Laughlin’s original assertion (Laughlin, 1981); we found ARUs to allocate their effective AF range proportionally to the input statistics, thereby mitigating variations in the output distribution. Further details about the role of this mean response value on the stability of network dynamics are presented in Section 2.4.

**Figure 3.**
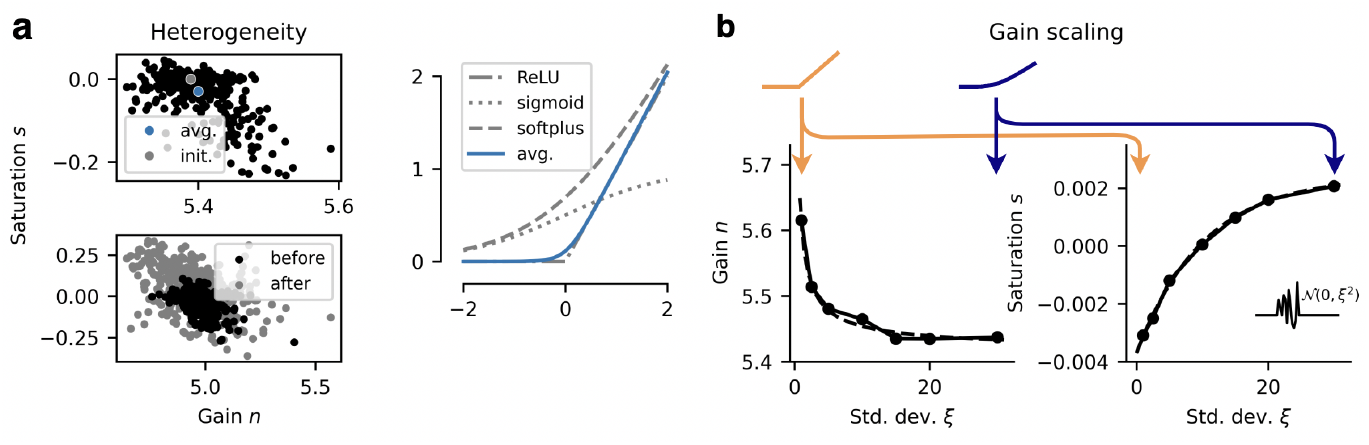
Steady-state coding mechanisms of ARUs. **(a)** Neuron-to-neuron heterogeneity in AFs. (Top) Learned activation parameters in RNN+*γ* (het.), from homogeneous initialization. (Bottom) Increase in heterogeneity before and after a 45*°* rotation is applied to the psMNIST digit, learned through backpropagation. (Right) The learned activation function shape corresponding to the average {*n, s*} parameters on the left (Top), shown along common AFs part of the *γ* AF family. **(b)** As a response to a noisy external drive of varying variance, the gain of the ARUs displays power-like decay (fit in dashed) as a function of the std.-dev. *ξ*, and the saturation displays exponential ramp-up (fit in dashed). Averaged over *N* = 3 seeds. Depictions of the associated *γ* AFs are included above, exagerated for visualization purposes (color follows colorbar).

#### ARUs as fractional integrators/differentiators

We find that ARUs exhibit neural integrator/differentiator behavior, which we investigate under the lens of fractional differentiation. Fractional order differentiation is a fundamental mechanism for efficient information processing in cortical neurons intimately intimately related to gain scaling (Anastasio, 1994; Lundstrom et al., 2008; Weber et al., 2019). It is defined by the product in Fourier domain *f* of the transformed signal with the operator *H*(*f*) = (2*iπf*)^*α*^. This is equivalent to a convolution in time domain. By varying the fractional order parameter *α* ∈ ℝ, one continuously interpolates between different order of differentiation of a signal. For *α* = 1, this operation is exactly your standard derivative; for *α* = −1, it is your standard integral. Both positive and negative orders are mathematically justified, observed in retinal neural populations (Kastner and Baccus, 2011, 2013), and carry meaning as a convolutional filter mechanism.

The response of the ARUs under different drives is mediated by the adaptation controller RNN, whose outputs are precisely the gain *n*_*t*_ and saturation *s*_*t*_. We observe prototypical onset {*n*_0_, *s*_0_} with exponential decay to steady-state values in the signals (Fig. 4**a**). To assess the impact of these rapidly modulated activation parameter signals on the ARU activity, we determine the order *α* of fractional filtering by minimizing, over *α*, the mean square error (MSE) between the fractionally de-filtered ARU activity and the original step perturbation. We find this methodology to recover with good fidelity the original signal (see e.g. Appendix Fig. 10), to settle on a sharp minimum, and to yield persistent results on all three random seeds and both tasks (see examples in Fig. 10-12). This indicates that the ARU controller effectively employs the precise fractional differentiation filter in Fourier domain observed in cortical neurons (Weber et al., 2019).

**Figure 4.**
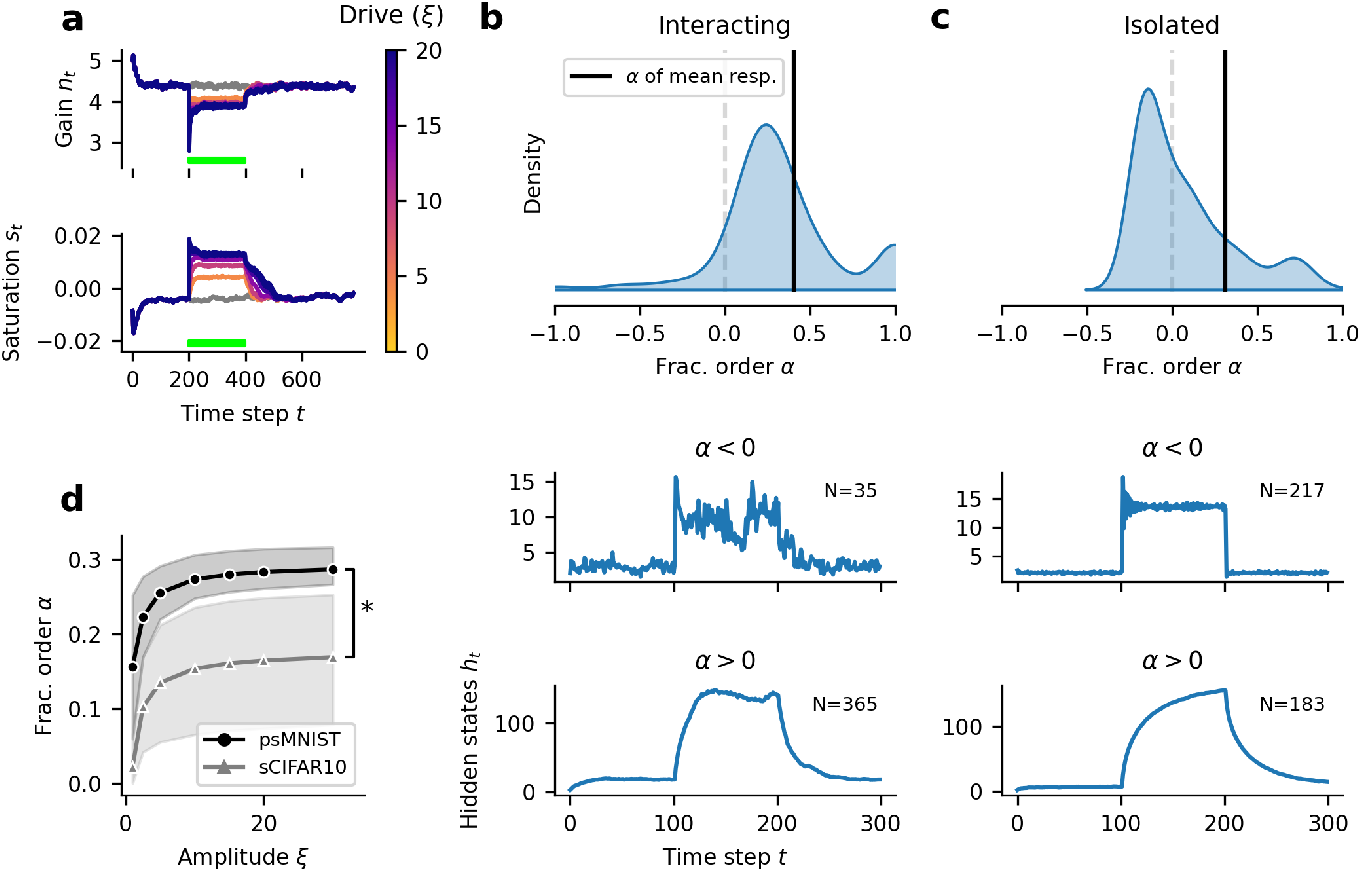
Fractional order integration and differentiation recovered by ARUs. **(a)** Gain *n*_*t*_ (top) and saturation *s*_*t*_ (bottom) signals during the processing of an input digit with an added step perturbation of varying amplitude *ξ*, applied during the green bar period. **(b, c)** Distribution, over neurons, of fractional order *α*. The order is established by minimizing the MSE between the fractional order differentiated signal of ARU activity and the step drive applied (*ξ* = 20 during *t* ∈ [100, 200)). If we apply the same procedure to the mean network activity, after averaging over neurons, we obtain the single estimate “*α* of mean resp” indicated by the black line. The two plots beneath represent the average activity for ARUs with the indicated fractional order *α*. **(b)** Fractional order *α* of ARUs with original fully connected recurrent dynamics. **(c)** Fractional order *α* of ARUs in isolation (diagonal recurrent weight matrix). **(d)** Fractional integration order, as a function of the step drive amplitude *ξ >* 0 and the task.

Specifically, we find heterogeneity in the fractional order across ARUs (Fig. 4**b**-top). Most units perform fractional order integration (*α >* 0, see Fig. 4**b**-bottom for average activity for those ARUs), on orders akin to sensitizing retina cells (Kastner and Baccus, 2011). To further elucidate this finding, we consider the same step-constant drives, now applied to single neurons in isolation, with trained weights. This is a setting conceptually closer to the original experiment by Ref. (Lundstrom et al., 2008), performed on slices of neocortical pyramidal neurons. In this “Isolated” setting, we observe the distribution in fractional order *α* over neurons to shift towards fractional differentiation (see Fig. 4**c**-top). Now we observe ARUs to fall on both sides of this filtering mechanisms, with more than half performing fractional order differentiation (Fig. 4**c**-middle) showing adaptive behvaior, and the others performing fractional order integration and showing sensitizing behavior (Fig. 4**c**-bottom). This provides additional evidence for the heterogeneity in neuronal types recovered in this framework.

Finally, taking a closer look at the exact fractional orders, we find that they depend on the task considered (Fig. 4**d**). For all *ξ* tested, we found the average fractional orders *α* to be different between the two tasks (*p <* 0.05 for *ξ* ≥ 1, *N* = 5 seeds, independent two-sample *t*-test). Furthermore, we did find the fractional order to depend on *ξ* (*p <* 0.05, *N* = 5, related two-sample *t*-test between *ξ* = 1 and *ξ* = 30), increasing for low values to a plateau for higher values.

### 2.4 Neural adaptation improves global network information propagation

So far, we have established that diversity and adaptive tuning of single neuron AFs improve an RNN’s performance on perceptual tasks, and enables enhanced robustness to noise. Furthermore, we showed that optimized adaptive dynamics are not only biologically more plausible, as they are implemented locally at each neuron, but they also implement dynamic coding mechanisms observed in real neurons (gain scaling, fractional differentiation). In this section, we further analyze the adaptive controller dynamics to uncover how ARU mechanisms emerge from end-to-end optimization, and probe what this suggests for the role of adaptation mechanisms in the brain.

We begin by focusing on the {*n*_*t*_, *s*_*t*_} trajectory, governing the AF of a single neuron when presented with a step-input. Recall that Fig. 4**a** shows such trajectories for different input amplitudes *ξ*. At stimulus onset, we observe an initial spike with *ξ*-dependent onset values we name {*n*_0_(*ξ*), *s*_0_(*ξ*)}. Following this is an exponential decay toward *ξ*-dependent steady state values {*n*_∞_(*ξ*), *s*_∞_(*ξ*)}. This transient decay is responsible for fractional differentiation and integration, while the the values {*n*_∞_(*ξ*), *s*_∞_(*ξ*)} dictate the magnitude of gain scaling, both reported in reported in §2.3. Fig. 5**a** shows the dynamics of the controller RNN whose output is the {*n*_*t*_, *s*_*t*_} trajectory, projected in the first two principal components of the controller’s hidden states (*N* = 50). Markers indicate steady states of the controller under constant input *ξ* following the transient decay, and lead to outputs {*n*_∞_(*ξ*), *s*_∞_(*ξ*)}. These clearly lie on a seemingly continuous 1-D manifold which encodes the amount of scaling the neuron’s AF receives under prolonged constant input. In what follows, we aim to uncover what coding advantages these adaptive dynamics afford, first at the single neuron level, then for networks.

**Figure 5.**
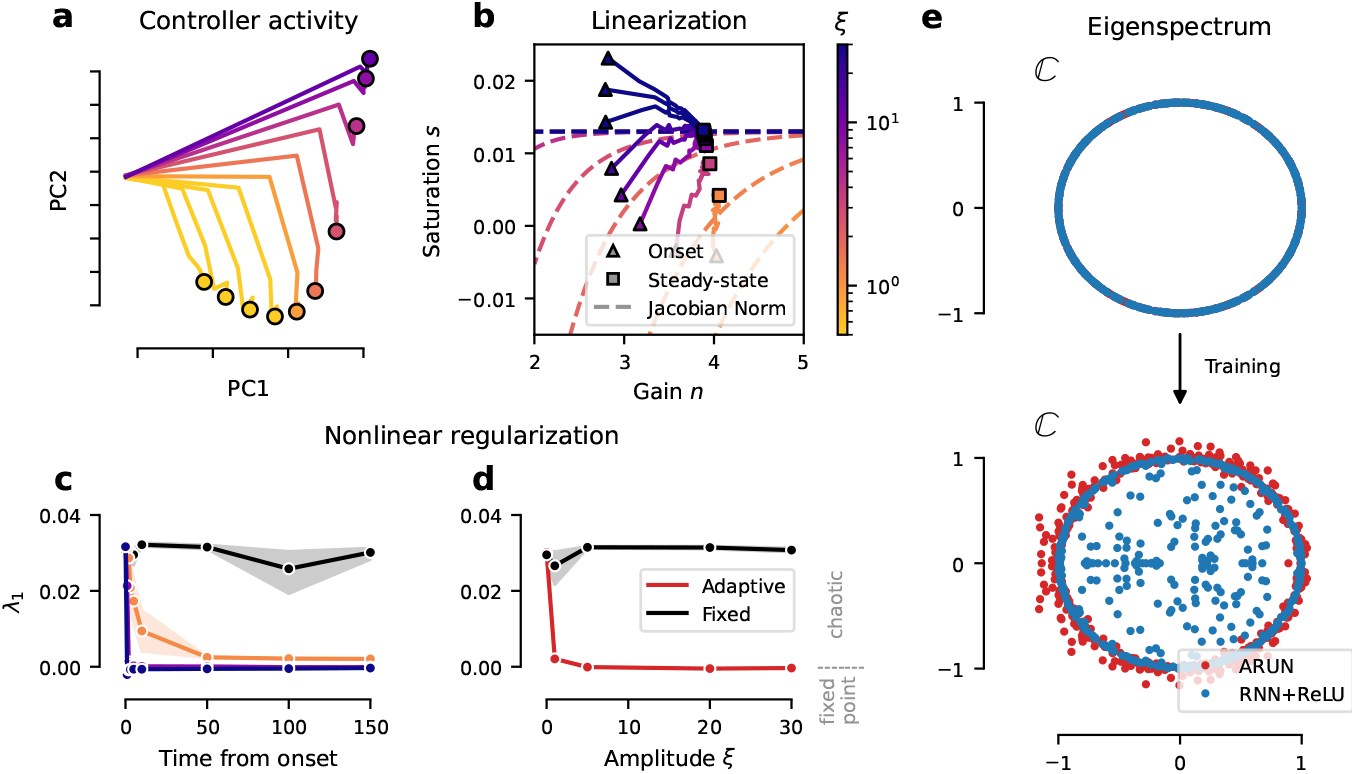
Dynamic regularization by ARU controllers. **(a)** Limit points in controller activity from constant input *ξ*. **(b) {***n*_*t*_, *s*_*t*_} trajectories during stimulation, with onset (triangle) and steady state (square), for each external drive *ξ*. Overlayed over jacobian *γ*^′^(*ξ*; *n, s*) = 1 *E* level curves in phase space (*E* = 0.01). Colorbar shared with (**a**-**c**). **(c-d)** Maximum Lyapunov Exponent (*λ*_1_) of an ARUN’s *main* RNN averaged over *n* = 10 draws from the stationary distribution (mean and 95% c.i. shown, details in Methods §5.4). The black line indicates the original *ξ* = 0 setting. (**c**) *λ*_1_ as a function of the chosen time-step 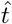 from the onset of the perturbation, for varying *ξ* (see colorbar in **b**). We take 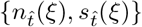 set by an ARU and treat them as unvarying in time (equivalent to RNN+*γ* static). (**d**) *λ*_1_ as a function of the step-drive amplitude *ξ*. We take the steady-state values *{n*_∞_(*ξ*), *s*_∞_(*ξ*)*}* (*ξ >* 0 for “Adaptive”, *ξ* = 0 for “Fixed”).

#### Adaptive activation reduces noise variance in single neurons

We now quantify the impact of the activation function *γ* and its parameters {*n, s*} on noise integration in single neurons. As a response to a general perturbation scalar *η* ∼ 𝒩 (*μ, σ*^2^), we seek to quantify the role of the parameters {*n, s*} in amplifying or reducing this noise. To this end, we present the following proposition.

##### Proposition 1.

*For unitary W*_*hh*_ *weight initialization, the variance explained along a vector* 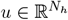 *as a response to a perturbation* ***η*** ∼ 𝒩 (***μ***, *σ*^2^*I*) *decays if and only if the parameters {****n***_*t*_, ***s***_*t*_*} satisfy*

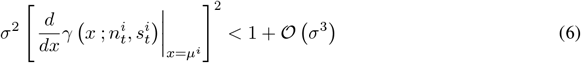

*for i* ∈ *{*1, …, *N*_*h*_*}*.

*Proof*. See proof in Supplementary Materials. (Appendix §C.2)

This result leverages known conditions on an RNN’s Jacobian under mild connectivity assumptions to ensure noise inputs are not amplified. For example, for a linear AF with slope *a*, we would require *σ* ≤ 1*/a* to avoid noise amplification. For a neuron with AF *γ*(*x*; *n, s*) and small *ϵ >* 0, the continuity of *γ* and the implicit function theorem guarantees that there exists a 1-D manifold in {*n, s*} -parameter space indicating a stability boundary. We can derive this manifold parametrized by the drive’s amplitude *ξ* by solving for 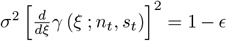. This path corresponds to the expected variation in activation parameters *{n, s}* as a function of *ξ* for the system to absorb, through the hidden-state dynamics, the injected noise by a margin *E*. This result requires linearization assumptions, see Appendix §C for further details.

Equipped with this result, we return to the ARU adaptive dynamics obtained by optimization. Panel **b** of Fig. 5 shows the same trajectories as Fig. 4**a**, plotted in the {*n, s*} -plane. Triangle markers indicate onset values {*n*_0_, *s*_0_} in response to step inputs, while square ones indicate steady states {*n*_∞_, *s*_∞_}. Dashed lines indicate the manifolds predicted by the above results, where color indicates corresponding input amplitude *ξ*. We can observe that for moderate to high *ξ*, our theory accurately predicts end states of {*n*_*t*_, *s*_*t*_} dynamics. The prediction degrades for low input amplitudes, a regime for which the main network can compensate noise by itself without neural adaptation (see Fig. 2). Overall, this indicates that in the presence of large perturbations (unseen in training), the adaptation-modulated gain-scaling operation described in Section 2.3 places neurons in a stable dynamic regime that avoids noise amplification.

#### Adaptation maintains network dynamics at the edge of chaos

Beyond single neuron mechanisms, we now turn to network-wide effects of adaptive units, and ask how adaptive units influence population dynamics. To better quantify this, we leverage Lyapunov Exponents, a measurement of average expansion and contraction of state space by a dynamical system. The sign of the largest exponent (*λ*_1_) is a good indicator of a system’s stability: chaotic systems have at least one positive exponent. A system with *λ*_1_ = 0 is said to be at the “edge of chaos”, with computational advantages we discuss in detail in the Discussion section. We approximate *λ*_1_ for RNNs as described in (Vogt et al., 2022) with, and without adaptation. We refer to Methods §5.4 for details on the computation, and Appendix D for details and a primer on the topic.

We find that for constant drives of distinct amplitude *ξ >* 0, adaptation mechanism actively steer the networks toward *λ*_1_ several orders of magnitude closer to 0 in comparison to non-adaptive RNNs (see Fig. 5**c-d**). This means that in situations where inputs are strong enough to destabilize dynamics in a non-adaptive RNN (thus leading to loss in performance), networks with ARUs remain more stable at the edge of chaos.

This mechanism is further showcased when observing the eigenspectra of optimized RNN connectivity matrices. In a standard non-adaptive RNN, connectivity matrices with eigenvalues whose magnitude are greater than one typically lead to chaotic dynamics, while smaller eigenvalues indicate stable dynamics. If RNNs were linear, the log eigenvalue norms directly indicate Lyapunov exponents. Therefore, it is widely known that initializing RNNs with unit norm eigenvalues leads to dynamics at the edge of chaos. One such initialization strategy is to initialize at the identity, or to select orthogonal matrices, also known as the Henaff initialisation (Henaff et al., 2016). In Fig. 5**e**, we compare optimized connectivity matrix eigenspectra of adaptive and non-adaptive RNNs with the same Henaff initialization and training procedure. What we observe is that adaptive RNNs eigenspectra have a magnitude well beyond the unit circle, in contrast to non-adaptive RNNs. Since we know from Lyapunov exponents that adaptive RNNs operate at the edge of chaos despite this connectivity, this indicates a form of “dynamic damping” provided by adaptation. As further explored in the discussion below, this suggests that adaptation at the single neuron level enables added expressivity by allowing a wider range of recurrent connectivity, while maintaining stability across dynamic regimes.

In sum, while our adaptation mechanism is local and independent across neurons, the combined effects actively balance global network dynamics in response to changes in input statistics and conserve stable computations.

## 3 Discussion

### Optimal information propagation through local regularization

Recurrent neural networks, whether biological or artificial, must balance *expressivity* and *stability* to implement complex computations. A network whose dynamics quickly converge to a single fixed point, for example, is quite stable but not expressive since very little mathematical transformations of inputs can take place (e.g. integration, amplification). In contrast, a network operating in a chaotic regime is expressive but unstable: its dynamics are rich but tiny perturbations lead to widely contrasting outcomes. The balance between these two requirements has been identified in several contexts, and is often referred to as the “edge of chaos” (Bertschinger and Natschläger, 2004). Dynamics close to the transition point between stable and chaotic dynamics offer optimal information propagation over long timescales, as well as rich transformations (Bertschinger and Natschläger, 2004; Legenstein and Maass, 2007; Boedecker et al., 2011). This regime has also been shown to be important in deep and recurrent artificial neural networks (Poole et al., 2016). Indeed, a rich theory of how large networks learn and implement computations shows that expressivity is maximized when dynamics are close to chaotic.

In machine learning, several strategies have been developed to ensure efficient training of artificial neural networks. Much of these strategies rely on global knowledge of networks (Pascanu et al., 2012), and global interventions on connectivity (Arjovsky et al., 2016; Le et al., 2015; Henaff et al., 2016; Lezcano-Casado and Martínez-Rubio, 2019; Kerg et al., 2019). For instance during training, batch normalization has proven surprisingly efficient at enforcing certain dynamical regimes. Gating units such as those found in LSTMs and GRUs have also been shown to help stabilize dynamics (see Krishnamurthy et al. (2022) for a review of the role of multiplicative gating). However, these processes are not biologically plausible as they are inherently non-local, requiring knowledge across space and/or time. While useful in AI/ML, these mechanisms offer little insight into how the brain might maintain dynamics at the edge of chaos under a variety of input statistics and distributed changes in connectivity throughout learning. Our results suggest that diverse and adaptive single neuron properties likely play such a role. Indeed, we demonstrate that ARUs offer a form of biologically plausible dynamic regularization, dampening the system in the presence of changing input statistics in such a way to promote network-level optimality. In our experiments, these mechanisms emerge from end-to-end optimization and bear striking similarity to well-studied biological neuron properties, suggesting adaptive neural dynamics could have evolved to maintain networks in optimal states.

### Deep learning and backpropagation as a framework to uncover biologically realistic optimal solutions

In this study, we train network models using gradient descent via backpropagation (through time, BPTT) of errors (Goodfellow et al., 2016), an algorithm that computes error gradients at once for all network parameters (including adaptation controllers). It is unlikely that the brain makes use of backpropagation exactly as it is implemented for artificial network optimization, thanks to the non-locality of credit assignment arising when computing gradients. Importantly, we reiterate that our experiments do not model biological learning, but rather leverages optimality of learned solutions to study emergent mechanisms. Nevertheless, an important question is to ask if optimal solutions found by backpropagation have any reasonable correspondence to those found by evolution and biological learning. We argue that the requirement of stable and expressive information propagation in time is a central commonality shared by optimized artificial and biological networks.

When training RNNs with BPTT, a central challenge is the vanishing and exploding gradient problem (Pascanu et al., 2012; Bengio et al., 1994). Here, gradient norms tend to either grow or vanish exponentially with time steps, leading to problems when conducting gradient descent. For an RNN whose dynamics are given by *h*_*t*+1_ = *F* (*h*_*t*_, *x*_*t*_), the main culprit for exponential growth and decay when computing error gradients comes from a term involving long products of Jacobian matrices evaluated along trajectories:

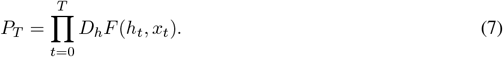

For RNNs following Eq. (1), *D*_*h*_*F* (*h, x*) = diag (*γ*^*1*^(*W*_*hh*_*h* + *W*_*xh*_*x*; *n, s*))^⊤^ *W*_*hh*_. Given idealized nonlinearity *γ* (e.g. ReLU) it should be clear that requiring eigenvalues of *W*_*hh*_ to be close to unit norm mitigates vanishing or explosion of *P*_*T*_. Crucially, we note that Eq. (7) is precisely the variational equation of a dynamical system (see (Vogt et al., 2022)), and that Lyapunov exponents are given by the time-averaged logarithm of spectral quantities of *P*_*T*_ when *T* → ∞.

Therefore, the stability of backward gradient propagation and of forward information propagation are two sides of the same coin. Forward dynamics at the edge of chaos, with Lyapunov exponents close to zero, correspond exactly to a regime where gradient norms avoid exponential growth or decay. In other words, a network operating at the edge of chaos forward in time also propagates error gradients backward in time efficiently. Thus, networks optimized via BPTT, which naturally promotes stable gradient propagation (Goodfellow et al., 2016), will be pushed to operate at the edge of chaos in forward time. If we assume the brain evolved to operate at the edge of chaos thanks to selective preference for neural circuits that propagate information over long timescales (a prevailing possibility with several experimental confounds, see e.g. (Bertschinger and Natschläger, 2004; Legenstein and Maass, 2007; Boedecker et al., 2011)), then there should be an important overlap between biological and artificial solutions for dynamic stability. As such, despite BPTT not being biologically plausible as a learning mechanism, we argue that ingredients contributing to stable information propagation that emerge from BPTT, such as adaptive mechanisms, are likely consistent with brain evolutionary pressures toward stable and expressive information propagation. We refer the reader to §C.5 for a detailed analysis of gradient propagation in our model. There, we compare gradient norms as they propagate throughout training and found that adaptive networks maintain stable gradient norm better than non-adaptive RNNs trained in the exact same way.

Finally, we note that we did not focus on the issue of multiple optimization timescales. In a more strict comparison to neuron dynamics and neural circuits in the brain, ARU controllers would have been optimized over evolutionary timescales, while the main RNN parameters, representing synaptic connections, over the lifespan of an animal. We did try a limited number of experiments, for example by fixing one while learning the other and vice versa (see Appendix §C.4), and did not see any significant differences in results. A different methodology, borrowing from deep learning frameworks like meta-learning, could allow for a more adequate consideration of adaptive mechanisms as a product of evolution-like pressures (see e.g. Sandler et al. (2021)). Such a more thorough investigation of the impact of learning timescales on solutions is outside of the scope of this paper, but is a fascinating direction of future work to disentangle evolution and learning pressures.

## 4 Conclusion

In this work, we sought to investigate goal-driven learning pressures from the system-level onto dynamic coding mechanisms at the single-neuron level. We do so by introducing *adaptive recurrent units*, allowing for online AF control from a novel parametric family. Our main findings are threefold: (1) Diverse and adaptive activation functions considerably improve computational performance of networks while also helping mitigate previously unobserved changes in input statistics during a task, thus improving out-of-distribution generalization. (2) System-level learning pressures drive biologically plausible adaptation strategies, namely activation function having biologically realistic configurations and more importantly, the implementation of biological SFA mechanisms such as gain scaling and fractional differentiation. (3) Finally we find that adaptation acts as a dynamic regularizer allowing recurrent neural networks to remain in a dynamic regime closer to the edge of chaos where forward and backward information propagation is optimal. These findings are supported by detailed numerical experiments and analytically derived bounds for information propagation in networks. We discuss how ARU adaptation can effectively implement a number of methods often used in deep learning to ensure good network expressivity and stability, including regularization and normalization. In contrast to these methods which require global, biologically unrealistic network quantities, ARU adaptation is local to each neuron and is consistent with known physiology. Taken together, our results support that neural diversity and adaptation serves a crucial role in goal-oriented network optimization, which suggests a coordinated and consistent optimality across scales linking brain circuits and single neurons.

## 5 Methods

### 5.1 Tasks

#### psMNIST

(Permuted Sequential MNIST). This task, first proposed by Le et al. (2015), focuses on learning to classify hand-written digits from the MNIST dataset (Deng, 2012) by presenting the pixels sequentially. This requires an accumulation of information over long timescales. Furthermore, we apply a fixed random permutation to the pixels; this reduces the time-step to time-step correlation and thus makes the task harder.

#### sCIFAR10

(Sequential CIFAR10). This second task is fundamentally similar to the previous MNIST one, where here the goal is to classify CIFAR10 (Krizhevsky et al., 2009) images. The network is shown (grayscaled) images of real world objects one pixel at the time and has to determine to which one of the 10 classes the image belongs. Now because the images are of objects in a variety of settings and not simply of digits, this constitutes a significantly harder task than psMNIST.

### 5.2 Network architecture details

For the RNN+*γ* and ARUN networks, we use a network size of *N*_*h*_ = 400 hidden-states. In the ARUN, the adaptation controller is chosen to operate with a smaller network, that we set to *N*_*g*_ = 50. With respect to a standard RNN, like the RNN+ReLU model, a heterogeneously optimized AF introduces 2 · *N*_*h*_ new parameters, and an adaptive controller RNN introduces (*N*_*g*_)^2^ + 2 · *N*_*g*_ parameters. We control the number of hidden-states in the other LSTM and GRU architectures to provide a similar (within 2%) number of parameters across all architectures.

We use linear encoder networks for the psMNIST task. For the CIFAR10 task, we resort to convolutional neural network (CNN) encoding schemes. More precisely we use a CNN with 4 convolutional layers and max pooling after the 2^nd^ and 4^th^ conv. layers. All convolutional layers are directly preceded by batch normalization and an element-wise application of the ReLU activation function and all layers have 3 × 3 sized kernels as well as padding and stride of 1 in both directions. The number of channels in each convolutional layer is 32, 64, 128 and 256 in order from input to output. In both cases max pooling is used over 2 × 2 sized regions with a stride of 2.

### 5.3 Modeling and training details

The vector of all trainable parameters is denoted Θ, and the parameters are updated via gradient descent using backpropagation through time (BPTT), with the matrix *W*_*rec*_ initialized using a random orthogonal scheme (Henaff et al., 2016). Independently of the task, we used Cross-entropy loss as our loss function and the Adam (Kingma and Ba, 2015) optimizer. We experimented with the RMSprop optimizer (introduced in Hinton et al. (2012), first used in Graves (2013)) with smoothing constant *α* = 0.99 and no weight decay, which yielded similar results.We trained the networks for 100 epochs. We investigated different learning rates (LR ∈ *{*10^−3^, 10^−4^, 10^−5^, 10^−6^*}*), and settled on 10^−4^.

#### Activation parameters

We first consider an initialization grid *N* × *S*, where *N* = {1.0} ⋃ {1.25*k* : 1 ≤ *k* ≤ 16} and *S* = {0.0, 0.25, 0.5, 0.75, 1.0} such that |*N*| = 17 and |*S*| = 5. For each pair of parameters 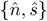, *ŝ* on the grid, we then evaluate the test accuracy on held-out digits for the psMNIST task with RNN+*γ* networks with *γ*(·; 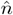, *ŝ*) AF. Here we consider only the static homogeneous setting. We plot in Appendix Fig. 6**c-d** respectively the performance landscape for a fixed static activation, and the evolution of the parameters *n, s* as they are updated with gradient descent alongside training. The resulting performance landscapes are similar to what would be expected from the Jacobian-norm and Maximum Lyapunov Exponent *λ*_1_; unit-norm jacobian, and the edge of chaos, is favorable. More details in Appendix §B.2. Therefore, we set our initialization prior to be *n* ∼ *N* (5, 2^2^*I*) and *s* = 0 for all further analysis.

**Figure 6.**
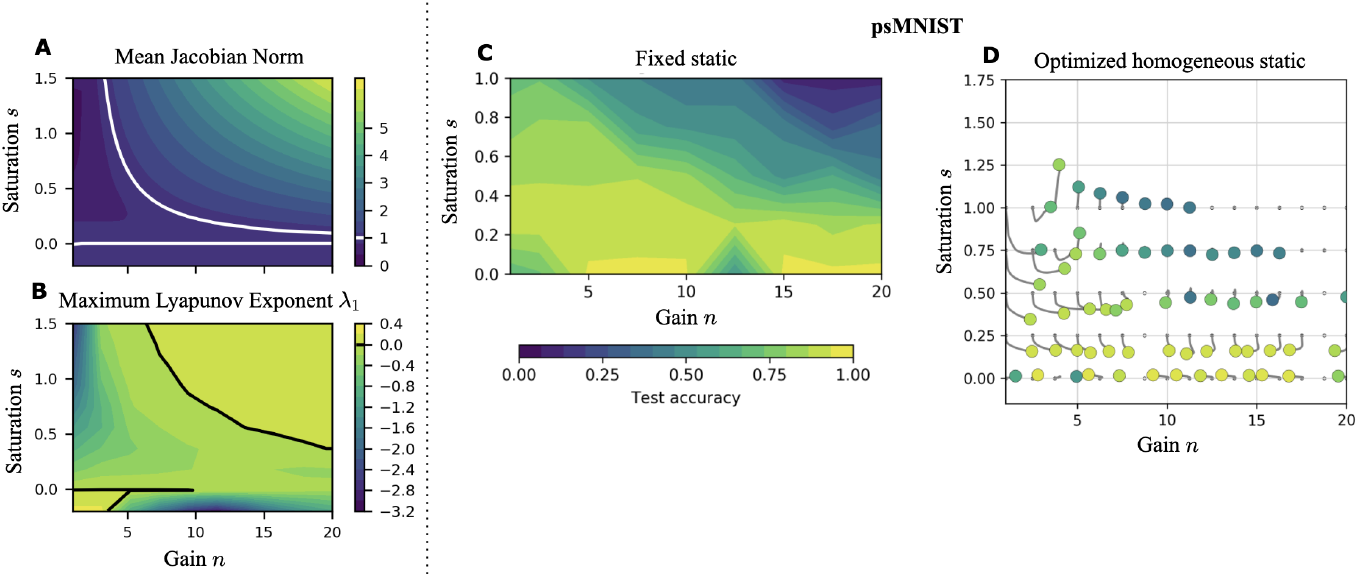
**A-B** Task independent stability metrics in activation parameter space. **C-D** Test accuracy in activation parameter space for the psMNIST task under two different learning scenarios.

#### Pytorch autograd implementation of gamma

We implement *γ*(*x*; *n, s*) as a Pytorch autograd Function with corresponding Pytorch Module.

To allow for activation function adaptation, we further include the activation parameters in the backpropagation algorithm. We do so by defining the gradient of *γ* with respect to the input and parameters. We can rewrite *γ*(*x*; *n, s*) as :

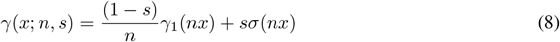

where *σ*(*x*) is the sigmoid activation function. With this notation, the partial derivatives of *γ* with respect to *x* (or total derivative), *n* and *s* are

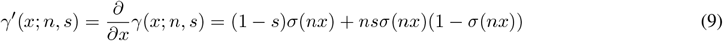

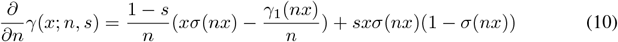

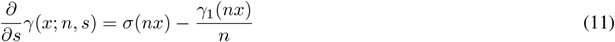

### 5.4 Evaluation methods and metrics

To assess how the activation gain and saturation influence on signal propagation, we use three quantities:

#### Jacobian norm

The main mechanism leading to the well studied problem of exploding & vanishing gradients in backpropagation and BPTT happens when products of Jacobian matrices explode or vanish (Pascanu et al., 2012; Bengio et al., 1994). We average the *L*^2^-norm of the derivative of Eq. (1) with respect to ***h***_***t*−1**_ ∼ 𝒰 (−5, 5). A mean **Jacobian norm (JN)** that is greater/less than one leads to exploding/vanishing gradients, respectively. An issue with this approximation is that the true mean depends on ***h***_***t***_’s invariant distribution, which changes with (*n, s*).

#### Lyapunov Exponents

Developed in Dynamical Systems theory, Lyapunov exponents measure the exponential rate of expansion/contraction of state space along iterates. Let us define *F* : *X* → *X* to be a continuously differentiable function, and consider the discrete dynamical system (*F, X, T*) defined by

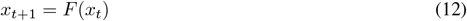

for all *t* ∈ *T*, where *X* is the phase space, and *T* the time range. Let *x*_0_, *w* ∈ *X*, define

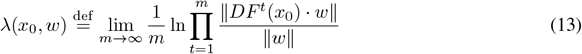

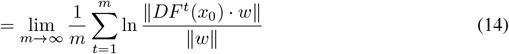

Note that once *x*_0_ and *w* have been fixed, the quantity *λ*(*x*_0_, *w*) is intrinsic to the discrete dynamical system defined by *x*_*t*+1_ = *F* (*x*_*t*_). We call *λ*(*x*_0_, *w*) a **Lyapunov exponent** of the mentioned dynamical system. As outlined in Appendix §D.2, it can be shown that under regularity conditions on the map *DF*^*t*^(·), the Lyapunov exponents effectively do not depend in the perturbation vector *w*, hence we consider *λ*(*x*_0_) only. Intuitively, Lyapunov exponents are topological quantities intrinsic to the dynamical system that describe the average amount of instability along infinite time horizons. Now, a randomly chosen vector, has a non-zero projection in the direction of the **Maximal Lyapunov exponent (MLE)** with probability 1, and thus over time the effect of the other Lyapunov exponents will become negligible. This motivates taking the MLE as a way of measuring the overall amount of stability or instability of a dynamical system. (see Appendix §D for a LE primer). As an asymptotic quantity, the MLE has been used to quantify ANN stability and expressivity (Pennington et al., 2018; Poole et al., 2016). The MLE gives a better measurement of stability than the Jacobian norm above, although it requires more effort to approximate. A positive MLE indicates chaotic dynamics and can lead to exploding gradients, while a negative MLE leads to vanishing ones.

We numerically compute the Maximal Lyapunov Exponent *λ*_1_(***h***_0_) for our RNN systems (eq. 1) using a QR algorithm (as motivated in Appendix D.3). The computation relies on the recurrent matrix *W*_*hh*_, the activation function *γ*(·; ***n, s***), and is a function of the initial condition ***h***_0_. We average the value of *λ*_1_ across i.i.d. ***h***_0_ draws from the stationary distribution *π*(*ξ*) of the process under a step-drive perturbation of amplitude *ξ* ≥ 0. For *ξ* = 0, this is the perturbation-less setting. If one consider 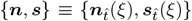 as the activation parameters setting by the ARUN at time 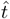 after onset under a drive *ξ* (as in Fig. 5**c-d**) this effectively makes *λ*_1_ a function of only 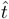 and *ξ* for fixed *W*_*hh*_.

#### Order of fractional order filtering

We investigate ARU activity under the lens of fractional differentiation (Anastasio, 1994; Lundstrom et al., 2008; Weber et al., 2019). The mechanism is defined implicitly by the product in Fourier domain *f* of the transformed signal with the operator *H*_*α*_(*f*) = (2*iπf*)^*α*^

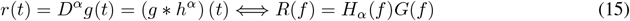

or can be thought of as a three step process,

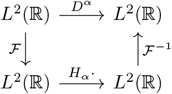

first transforming the signal *r* ∈ *L*^2^ in Fourier domain *R*(*f*), followed by a point-wise multiplication with *H*_*α*_(*f*), before finally performing an inverse Fourier transform back to the time domain. This procedure is exactly how we compute a fractional order differentiated signal, using the Discrete Fourier transform `numpy.fft.fft` (and associated `fft.fftfreq` and `fft.ifft` methods) from the Numpy (Harris et al., 2020) library.

In the specific case of determining *α* for ARUs, we operate under the hypothesis that the ARUs perform fractional order filtering of an input step signal *s*(*t*), such that their activity *r*(*t*) = [*D*^*α*^*s*](*t*). Hence we determine the order *α* of fractional filtering by minimizing, over *α*, the mean square error (MSE) between the fractionally de-filtered ARU activity and the original step perturbation. Specifically for *r*(*t*) the signal in time *t* in response to a step drive *s*(*t*), we determine 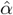 as

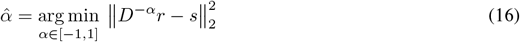

### 5.5 Network perturbations and task variations

To test the adaptive capabilities of our model and to compare it with conventional RNNs, we consider two different ways in which these external inputs may be perturbed:

1. **Variable contrast:** transforming the inputs *x*_*t*_ (*Wx*_*t*_ + *b*). A brightness factor from a randomly sampled sinusoidal curve may multiplies the *x*_*t*_ input at each time-step *t* (Figure 2). These transformed inputs are then encoded by the same linear module *W*_*xh*_.
2. **Perturbed:** applying an external drive directly to the neurons (*Wx*_*t*_ + *b*). Taking inspiration from optogenetic stimulation, we inject a scalar external drive *ξ* ∈ ℝ directly to the neuron (e.g. for ARUN, we add +*ξ* in equation (3)), before the activation function is applied. This perturbation may either be a non-random scalar (Figure 2) or noisy.

## Supplementary Material for

### A Experimental details

#### Task independent stability metrics

Figure 6 shows the task-independent stability metrics of JN and MLE for a range of (*n,s*) values (fixed across neurons). We see that the activation shape influences Jacobian norms, thus confirming that it will play an important role during training. Consistent with the average gradient norm, the MLE reports distinct (*n, s*)-regions of stability for random networks. In some cases, expansion and contractions can be useful for computations, and we further use these measurements to study training dynamics.

### B Performance: supplemental

#### B.1 Comparison between homogeneous and heterogeneous activation functions

Within the optimized learning framework, where the activation parameters {*n, s*} are learned by gradient descent of the loss, one could decide to enforce the activation function to be shared by all neurons (*homogeneous*) or vary neuron-to-neuron (*heterogeneous*). We incorporate this diversity by setting scalar {*n, s*} parameters in the homogeneous case, and by setting vectors ***n, s*** ∈ ℝ^*N*^ in the heterogeneous case for *N* neurons, such that the activation function of neuron *i* is set by ***n***^*i*^, ***s***^*i*^.

In terms of performance for the psMNIST task, we found that learning heterogeneous activation provided a slight increase but no significant advantages over the already well performing optimized homogeneous setting. On the gsCIFAR10 task the same conclusions hold, the heterogeneous RNN+*γ* performs as well if not slightly better than the homogeneous RNN+*γ* (see Figure 2), however the increase in performance is again not statistically significant.

#### B.2 Further details on learning differences and performance in the static setting

As expected, we find a strong correlation between the norm of the Jacobian in parameter space which is task-independent (Fig. 6**A**) and the performance landscapes for each task (see Fig. 6**C**). Interestingly, regions in space (*n, s*) with poor performance are all associated with an exploding gradient, not a vanishing gradient. Networks whose activation functions have activation parameters in a neighborhood of { (*n, s*) : ‖*γ*^*1*^(*x*; *n, s*) ‖ = 1} present optimal performance, on all the tasks. On the one hand, this further emphasizes the performance of ReLU (see (Glorot et al., 2011)) as part of this (*n, s*)-neighborhood. However, as we show in Fig. 6**C**-**D**, traditional nonlinearities (including ReLU) are outperformed by the considerably different activation functions arising in the different scenarios of end-to-end learning. This result highlights that non trivial combinations of parameters may also achieve optimal performance while allowing for much more complex nonlinear transformations than ReLU.

**Figure 7.**
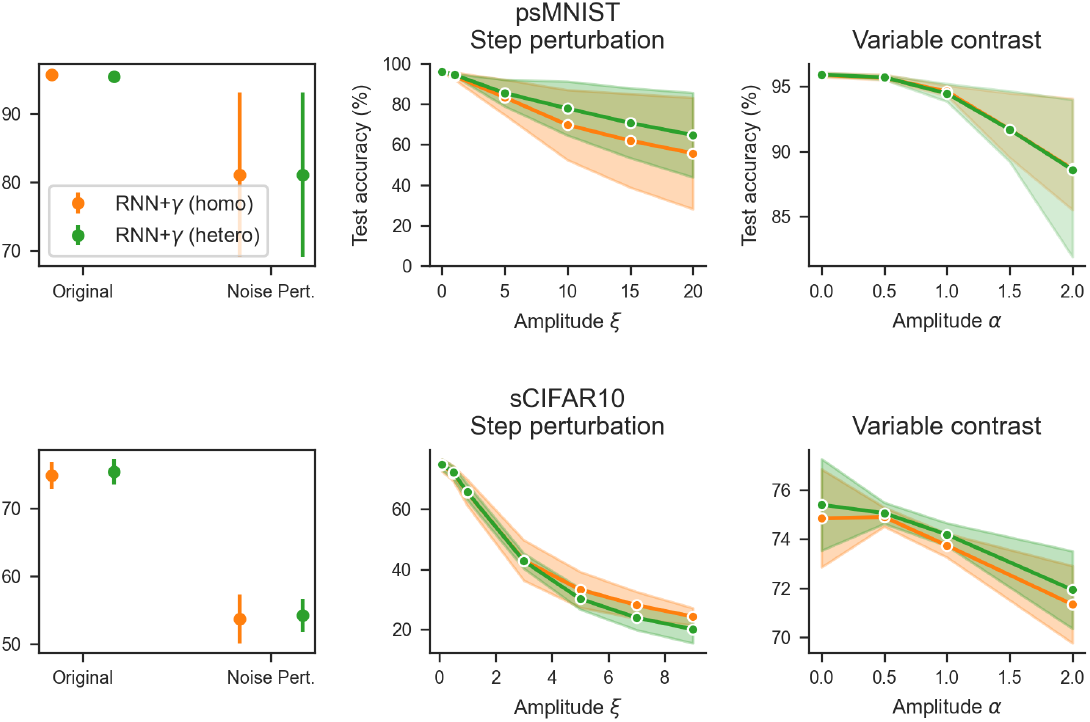
Comparison between homogeneous and heterogeneous activation functions. Labels and perturbation details follow Figure 2.

#### B.3 Learned adaptation offers transfer learning advantages

In neuroscience, the term adaptation is mostly used to describe processes that occur on short timescales and at a neuron level which have been shown to account for changes in stimulus statistics (Weber et al., 2019). This mechanism is naturally linked to the concept of transfer learning in AI where one seeks systems where minimal changes in parameters allow adaptation from learned tasks to novel ones. To see if changes in single neurons activation could offer transfer advantages in ANNs, we design a novel task using the psMNIST test data set where the images are rotated by 45°. The goal is for a trained network to adapt to this change in input structure by only changing its activation function parameters. To evaluate this, we split rotated images into training and test sets, each containing approximately 5k images and the same number of images per digit. We then briefly retrain heterogeneous activation parameters (*n*_*i*_, *s*_*i*_) on this rotated data set using the heterogeneous adaptation scenario. For initialization, we take the parameters (including the (*n*_*i*_, *s*_*i*_)’s) that resulted from training with normal images, also under the heterogeneous adaptation scenario. Before retraining, the networks achieved an accuracy of 94% on the original data set, this fell to 42% after rotation. Retraining (*n, s*) allowed the networks to recover classification accuracy up to 56%. This shows that simply allowing the activation functions to adapt can recover over a quarter of lost performance (over 25%). An example of the variation of (*n, s*) trajectories after retraining is showed in Fig. 3**a** (bottom).

**Figure 8.**
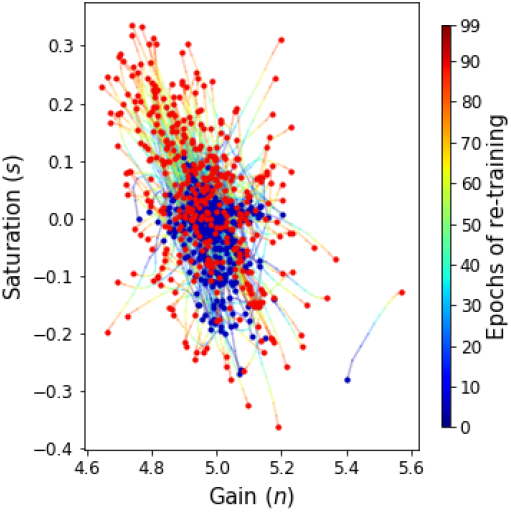
Trajectories of the activation parameters during retraining on the modified MNIST images.

Like in Fig. 3**a** (top), the cloud of (*n, s*) parameters expands with respect to its initialization, suggesting that a diversification in activation function shapes is needed to adapt to the change in task.

Allowing for small changes in the activation functions of individual neurons helps to mitigate the loss in performance caused by drastic changes in network inputs. The following question naturally arises from these results: is it possible to leverage the advantages brought by adaptation in an online manner instead of relying on retraining a part of the network? Such a “dynamic” adaptation process, which allows the network’s activation functions to instantly change when presented with inputs of different statistics, would not only be less computationally expensive and faster but would also be more alike its natural counterpart. We further explore the idea of implementing rapid adaptation protocols for ANNs in the next section.

### C Adaptation: supplemental

#### C.1 Fractional differentiation

##### Activation function parameters

Further details on fractional order differentiation of the activation parameter signals, as opposed to the resulting hidden-states *h*_*t*_, is included in Figure 9.

**Figure 9.**
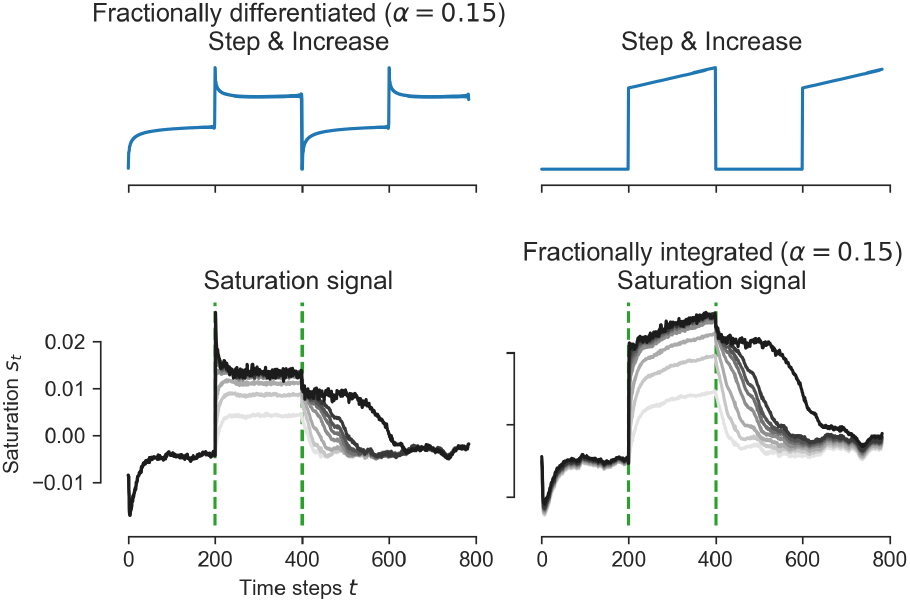
(**Top**) Graph of a step to linear-increase function (right), then fractional order (*α* = 0.15) differentiated (left). (**Bottom**) Saturation *s*_*t*_ as a function of the time (left), for varying external drives *ξ* ∈ [0, 30] with the usual range applied during a stimulation period framed by the two dashed green lines. See next Figure 10 for colorbar. (right) The saturation signals *s*_*t*_ fractionally integrated with *α* = 0.15 reveal step to linear increase signals during the stimulation period.

**Figure 10.**
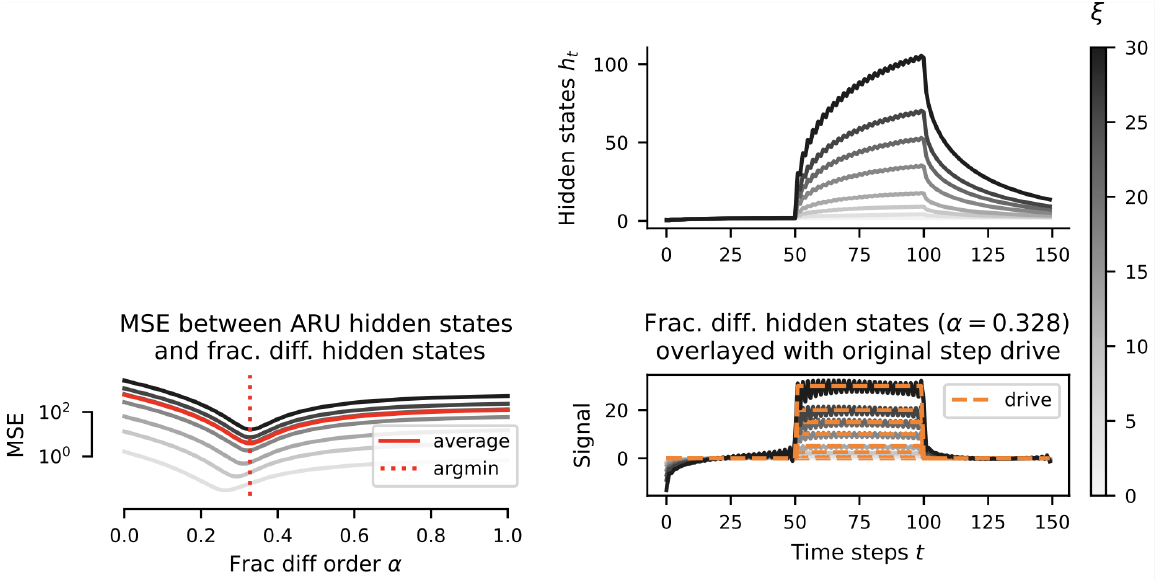
Task: psMNIST. Random seed #: 400. Colorbar applies to whole figure. (**top-right**) mean ARU hidden-states for non-interacting ARUs, just as main text’s setting. For other panels, see respective titles.

##### Determination of fractional order

The order *α* of fractional order differentiation was determined as the arg-min (over *α*) of the mean square error between the fractional *α*-order integrated signal and the precise step inputs that drove the network. See Figure 10, and more details on the methodology can be found in Methods §5.4. We observe that this minimum is sharp, and observe close correspondence between the fractional order integrated signal and the original step-drive. This analysis was consistent across tasks and random seeds (see examples in Figures 10, 11, 12).

**Figure 11.**
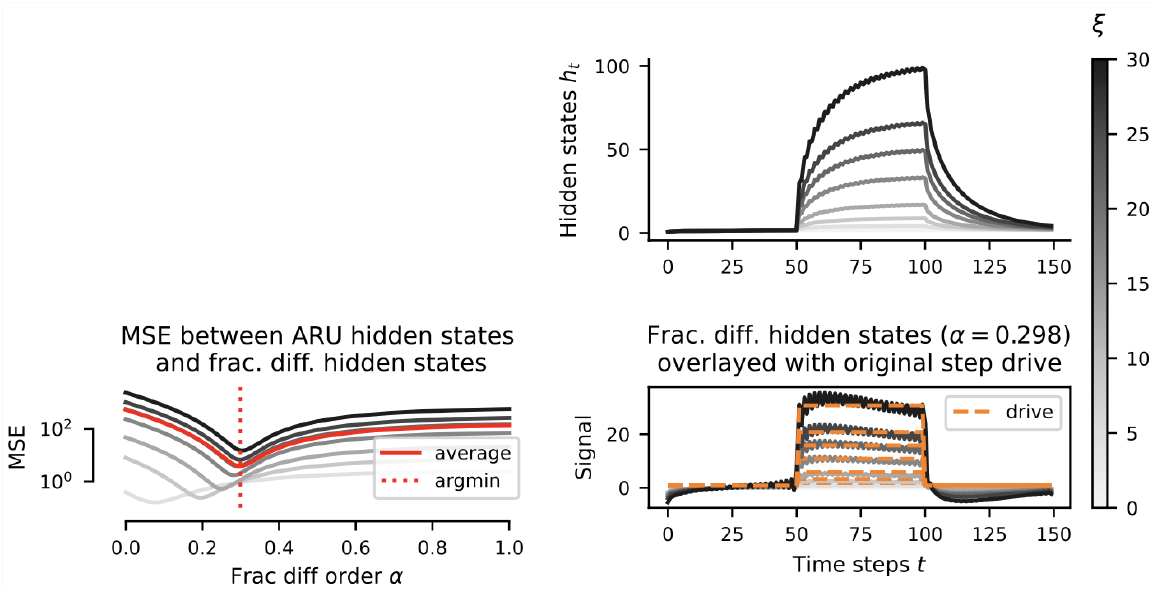
Task: psMNIST. Random seed #: 500.

**Figure 12.**
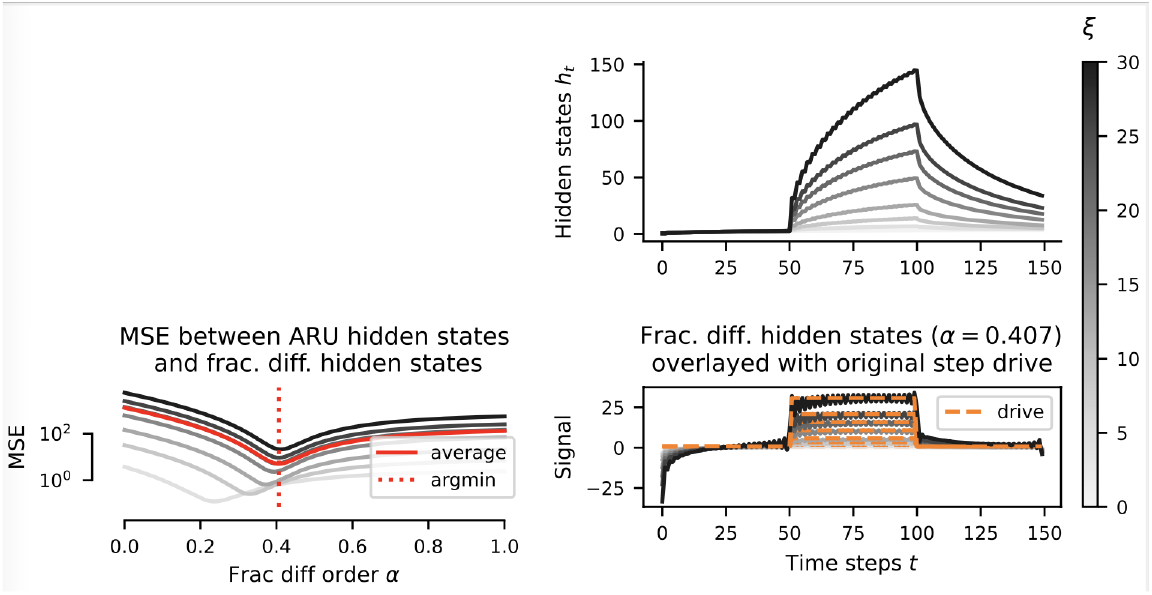
Task: sCIFAR10. Random seed #: 403.

#### C.2 Dynamic regularization

##### Proposition 1.

*For unitary W*_*hh*_ *weight initialization, the variance explained along a vector* 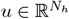 *as a response to a perturbation* ***η*** ∼ 𝒩 (***μ***, *σ*^2^*I*) *decays if and only if the parameters {****n***_*t*_, ***s***_*t*_*} satisfy*

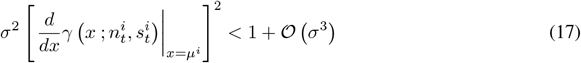

*for i* ∈ *{*1, …, *N*_*h*_*}*.

*Proof*. Consider some multivariate Gaussian noise *η* ∼ 𝒩 (*μ, σ*^2^*I*), injected in the dynamics

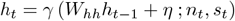

Now, the variance of this noise along a given vector 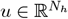 as it propagates through the dynamics is given by:

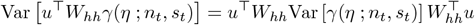

after one iteration. Since *η* is chosen such that *η*_*i*_ is independent of *η*_*j*_ for *i* ≠ *j*, i.e. Cov[*η*_*i*_, *η*_*j*_] = *σ*^2^*δ*_*ij*_, and *γ*(·) acts element-wise, we have that Cov [*γ*(*η*_*i*_), *γ*(*η*_*j*_)] = 0. As such,

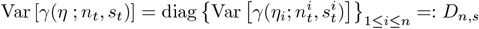

and

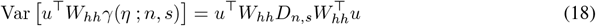

Using a first order Taylor expansion of *γ*(*x*; *n*_*t*_, *s*_*t*_) about the mean of *η*, we obtain an approximation of the variance

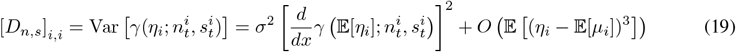

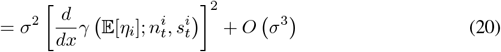

where the first term of the RHS can easily be evaluated directly (see SM equation (10) for a closed form expression of 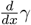). Also, under the initialisation schemes considered in our experiments, *W*_*hh*_ is unitary and as such 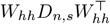 defines a normal matrix with eigenvalues exactly given by the entries of diagonal matrix *D*_*n,s*_. This gives the result.

This result reformulates known conditions on an RNN’s Jacobian and indicates that under mild connectivity assumptions, the left hand side of must remain smaller than one to avoid noise amplification. For example, for a linear AF with slope *a*, we would require *σ* ≤ 1*/a*.

#### Parameter evolution for noise integration

Let us for a moment restrict our attention to a single neuron, thus removing subscripts *i* and assuming scalar quantities. We note in passing that *σ* is non-zero even for scalar *ξ* as our formalization accounts for the linearly scaled inputs *x*_*t*_, which are distributed under the task input statistics. Now, consider the level set

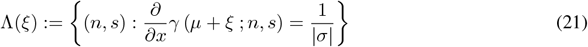

consisting of (*n, s*) values at the boundary of the region derived from Proposition 1 for a noise shifted by an external drive *ξ* ≥ 0 (un-perturbed if *ξ* = 0). As mentioned earlier this set corresponds to a manifold in (*n, s*) space, one that shifts as a function of *ξ* (see Fig. 5**b** for a visualization of these curves). Take 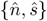, *ŝ* satisfying Prop. 1, and assume that there exists *E >* 0 for which *d*(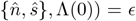, *ŝ*, Λ(0)) = *ϵ*. For noise robustness to be maintained in stimulated regimes, we have that the activation parameters *n*(*ξ*), *s*(*ξ*) should shift to stay within the region highlighted by Prop 1, i.e. *d*(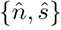, *ŝ*, Λ(*ξ*)) *>* 0 for all *ξ* ≥ 0. This is what we observe, see Fig. 5**b**. By fixing this distance and given an initial condition *n*_0_, *s*_0_, one can solve the above system to obtain a path *n*(*ξ*), *s*(*ξ*) in parameter space as a function of *ξ* (assuming continuous dependence on *ξ*). This path corresponds to the expected variation in activation parameters *n, s* as a function of *ξ* for the system to absorb, through the hidden-state dynamics, the injected noise by a margin *ϵ*.

Still, this does not account for particular behavior observed of an onset value {*n*_0_, *s*_0_} decreasing or increasing with an exponential time-constant to a steady-state value {*n*_∞_, *s*_∞_}, in a matter akin to spike frequency adaptation. Both onsets and steady-states satisfy the observations previously highlighted, but their–distinct– existence is unaccounted for. This sets a rich ground for future work.

**Figure 13.**
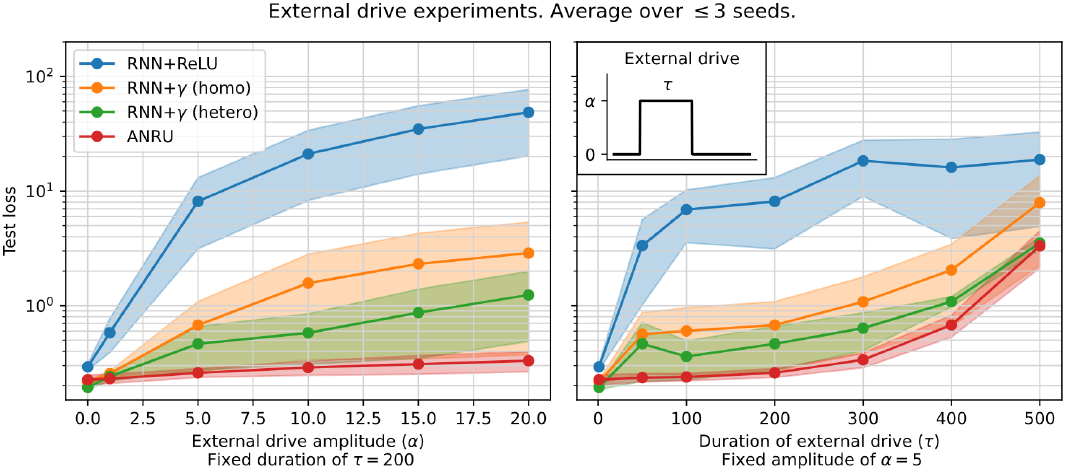
Sensitivity analysis for the step drive experiment. Lower is better. ARUN performs the best.

**Figure 14.**
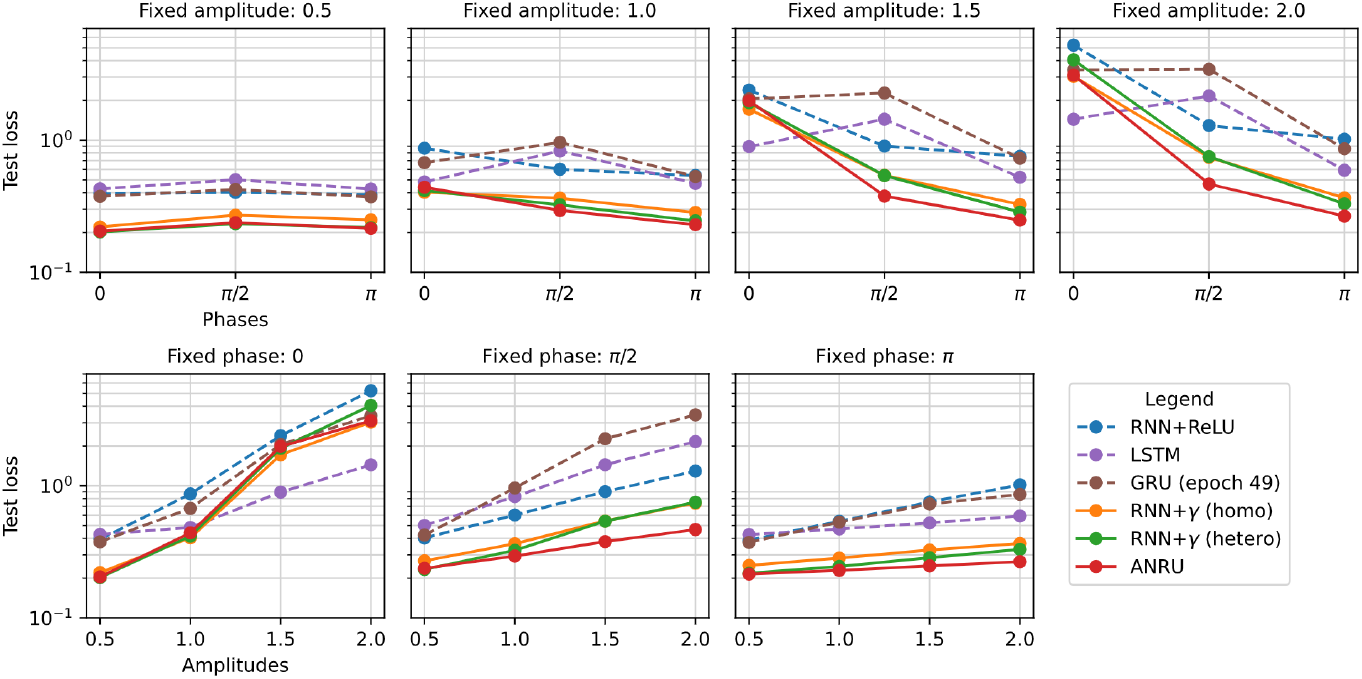
Sensitivity analysis for the Sinusoidal transformation on inputs, varying phase and amplitude alternatively. Lower is better. ARUN performs the best on average.

### C.3 Sensitivity analysis for the perturbations experiments

We refer to Figures 13-16.

### C.4 Testing the evolutionary plausibility of our adaptive units

The performance and robustness results presented in the main paper were obtained by randomly initializing and then simultaneosly training both the main RNN networks as well as the adaptive sub-networks. For our adaptive units to adequately model adaptation in biological neurons, the adaptive sub-units of each network should in principle be fixed when training the main RNN network. Indeed, in the brain, single neuron adaptation mechanisms have been developed over evolutionary timescales and are passed down through genetic information.

In this section we test our AURNs in a more biologically plausible setting, and see if the structure of the adaptive sub-network can be efficiently passed down from a network to another without affecting the network’s performance or robustness. To verify this we have tested the performance and robustness to the noise, step and sine data transformations of AURNs generated using two distinct initializing and training scenarios:

**Figure 15.**
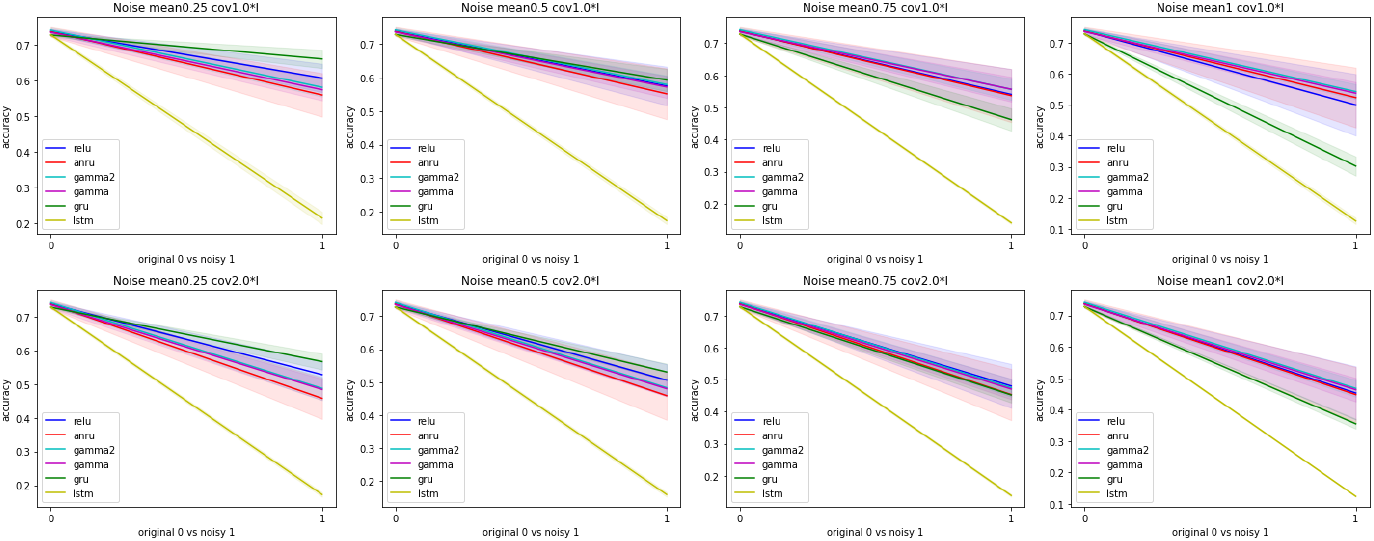
Sensitivity analysis for the noisy step drive experiment for the sCIFAR10 task. Higher is better.

**Figure 16.**
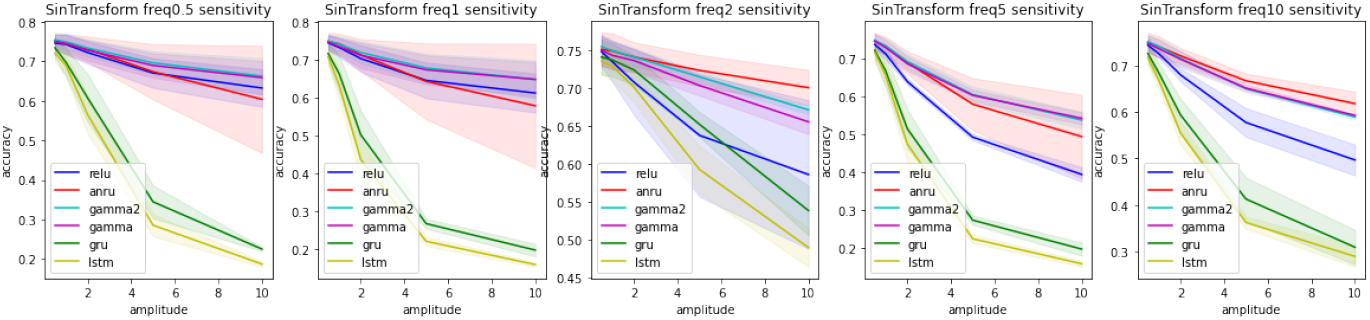
Sensitivity analysis for the sinusoidal transformation on inputs, varying the frequency, for the sCIFAR10 task. Higher is better.

- **Scenario 1:** Both the main and the adaptive RNNs are randomly initialized, using a specified random seed (here denoted *seed*_1_), and trained from scratch as previously described. All results from the paper are obtained with AURNs generated using this scenario.
- **Scenario 2:** The main RNN is randomly initialized using a specified random seed (*seed*_2_) while the adaptive sub-network is taken from a scenario 1 trained AURN with *seed*_1_ ≠ *seed*_2_. The main RNN is then trained using the same training procedure as in scenario 1 but the adaptive sub-network’s parameters are kept constant.

This was done for multiple random seeds of both the main RNN and the trained adaptive RNN, the results are shown in Fig. 17. We can see that the adaptation mechanisms previously learned with a specific main network can be used, as efficiently, by another main network without needing any re-training of the sub-network. The performance and robustness to different perturbations are, for all practical purposes, the same in both the setting where the main and the adaptive networks were trained simultaneously (scenario 1) and the setting where the adaptive sub-network was imported from a previously trained network and only the main network was trained (scenario 2).

**Figure 17.**
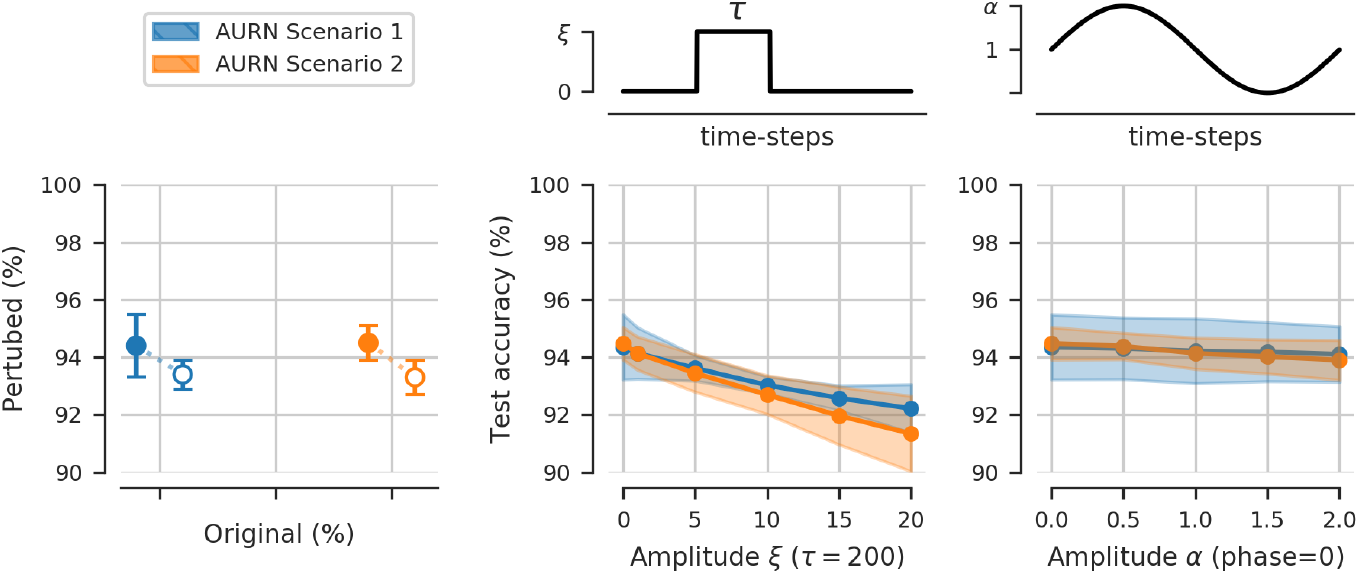
Performance on the psMNIST classification task and robustness to the noise perturbed, step drive transformed and sine transformed inputs. The mean and standard deviation across three different initializations are shown.

### C.5 Gradient contribution according to position in input sequence

The well documented vanishing and exploding gradients problems of RNNs prohibit effective training over long timescales. In particular, the gradient of the loss computed with the output of the network at time-step *t* + *δ* with respect to the hidden states of the network at time-step *t* either decays or increases exponentially with *δ*. In the vanishing case this makes the learning of long term dependencies impossible, while in the exploding case the entire training procedure is compromised. This phenomenon is linked to dynamic regimes in which an RNN operates, and is thus related to the leading Lyapunov exponent measurement described above (see Vogt et al. (2022); Poole et al. (2016) for more details).

We quantify the effects of learned neural adaptation on gradient propagation in RNNs by computing the gradient norms of the hidden-to-hidden weight matrix (*W*_*hh*_ in equations 1 and 3) on the psMNIST training set starting the gradient accumulation at different points in the input sequence. This was done for trained networks to take into consideration the learned adaptive behavior when considering the gradient propagation. In RNN+ReLU networks and to some lesser extend in RNN+*γ* heterogeneous networks the gradient norm increases monotonously with sequence length, as shown in logarithmic scale in Fig. 18. The earlier the accumulation of the gradients is started for these two network types the larger their norm is at the end of the input sequence when the loss is computed. In ARU networks however, after an initial increase the norms of the gradients actually decrease with sequence length. When the gradients are computed using the entire input sequences, the norm of the *W*_*hh*_ gradients in ARU networks is an order of magnitude smaller than in RNN+ReLU or in RNN+*γ* heterogeneous networks which promotes trainability and the stability of the gradient propagation during the training procedure. We also note that in ARU networks, elements (pixels) which are at the beginning of the input sequence and further away from the moment the loss is computed actually contribute more to the gradient of the weights when compared to later inputs. This is in stark contrast with gradient contribution in RNN+ReLU networks where the gradient contribution is monotonously increasing with the element’s position in the input sequence, early inputs contributing much less than later inputs to the gradient.

**Figure 18.**
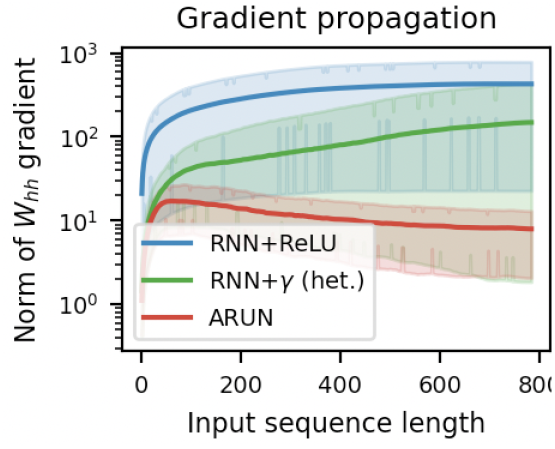
Frobenius norm of the hidden-to-hidden weight matrix *W*_*hh*_ gradient contribution of a given input element, or pixel, as a function of that element’s position in the input sequence. The sequences are of length 784 and elements closer to position 0 are closer to the beginning of the input sequences. The gradients are computed in trained RNN+ReLU, RNN+*γ* heterogeneous and ARU networks on the psMNIST training set. Mean and standard deviation across three random initialization are shown.

## D A primer on Lyapunov exponents

In this section we are first going to give a bit of theoretical background on Lyapunov exponents. Exponential explosion and vanishing of long products of Jacobian matrices is a long studied topic in dynamical systems theory, where an extensive amount of tools have been developed in order to understand these products. Thus one can hope to leverage these tools in order to better understand the exploding and vanishing gradient problem in the context of RNNs.

### D.1 Definition of Lyapunov exponents

Let *F* : *X* → *X* be a continuously differentiable function, and consider the discrete dynamical system (*F, X, T*) defined by

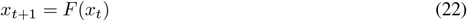

for all *t* ∈ *T*, where *X* is the phase space, and *T* the time range. We would like to gain an intuition for how trajectories of the mentioned dynamical system behave under small perturbations.

Let *x*_*t*_ and 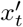 be two trajectories with initial conditions *x*_0_ and 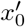, such that 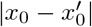 is sufficiently small. Defining 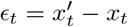, we get by the first order Taylor expansion

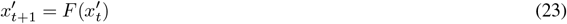

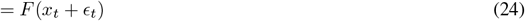

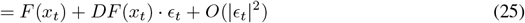

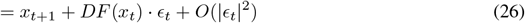

Substracting *x*_*t*+1_ both sides we get the variational equation

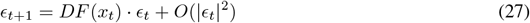

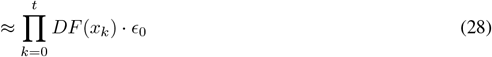

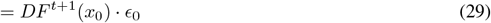

(Here *DF*^*t*+1^(*x*_0_) is an abuse of notation for the Jacobian of the (*t* + 1)-th iterate of *F*, evaluated at *x*_0_). Intuitively the ratio 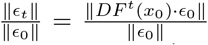 describes the expansion/contraction rate after *t* time steps if our initial perturbation was *E*_0_, which motivates the following definition:

Let *x*_0_, *w* ∈ *X*, define

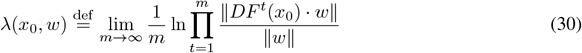

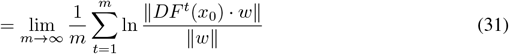

Thus *λ*(*x*_0_, *w*) measures the average rate of expansion/contraction over an infinite time horizon of the trajectory starting at *x*_0_, if it has been given an initial perturbation *w*. Note that once *x*_0_ and *w* have been fixed, the quantity *λ*(*x*_0_, *w*) is intrinsic to the discrete dynamical system defined by *x*_*t*+1_ = *F* (*x*_*t*_). We call *λ*(*x*_0_, *w*) a **Lyapunov exponent** of the mentioned dynamical system.

Since the Lyapunov exponents describe the the average rate of expansion/contraction for long products of Jacobian matrices, it doesn’t sound too surprising that they might provide an interesting perspective to study the exploding and vanishing gradient problem in RNNs. To give a complete picture of the analogy to RNNs, one can think of *x*_*t*_ as the hidden state at time *t*, and *F* can be seen as the function defined in the RNN cell. The only difference is that in RNNs we have inputs at every time steps, and thus the function *F* changes at every time step. This is the distinction between autonomous and non-autonomous dynamical systems, which is explained in more detail in the upcoming subsection D.4.

Finally, let us remark that the expression in the above definition of Lyapunov exponents is not always well defined. This will be the topic of the next subsection D.2, where we are presenting Oseledets theorem which gives exact conditions for when the above expression in well-defined.

### D.2 Oseledets theorem

As already stated, we bypassed the fact that the limit in the definition of *λ*(*x*_0_, *w*) might not actually exists. In fact this is the result of the well-known *Oseledets theorem*, but before stating the theorem let us point out a definition.

#### Definition.

A *cocycle* of an autonomous dynamical system (*F, X, T*) is a map *C* : *X* × *T* → ℝ^*n×n*^ satisfying:

- *C*(*x*_0_, 0) = Id
- *C*(*x*_0_, *t* + *s*) = *C*(*x*_*t*_, *s*)*C*(*x*_0_, *t*) for all *x*_0_ ∈ *X* and *s, t* ∈ *T*

#### Oseledets theorem

(sometimes referred to as Oseledets *multiplicative ergodic theorem*) Let *μ* be an ergodic invariant measure on *X*, and let *C* be a cocycle of a dynamical system (*F, X, T*) such that for each *t* ∈ *T*, the maps *x* ↦ log ‖*C*(*x, t*) ‖ and *x* ↦ log ‖*C*(*x, t*)^−1^‖ are *L*^1^-integrable with respect to *μ*. Then for *μ*-almost all *x* and each non-zero vector *w* ∈ ℝ^*n*^ the limit

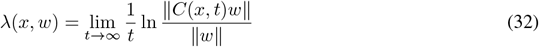

exists and assumes, depending on *w* but not on *x*, up to *n* different values, called the Lyapunov exponents (giving rise to a more general definition)

One can prove that the following matrix limit

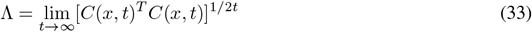

exists, is symmetric positive-definite and its log-eigenvalues are the Lyapunov exponents. We call Λ the *Oseledets matrix*.

In order to make this definition a little bit more intuitive, let us come back to our original situation, and note that the terms 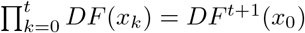 define a cocycle verifying the conditions of the theorem. Thus, in this case, the Lyapunov exponents are not only well defined, but there are up to *n* distinct ones of them, and they are the log-eigenvalues of the following Oseledets matrix:

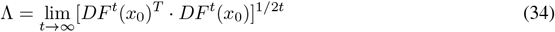

Let us now consider the singular value decomposition of *DF*^*t*^(*x*_0_),

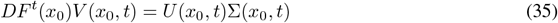

where Σ(*x*_0_, *t*) is a diagonal matrix composed of the singular values *σ*_1_(*x*_0_, *t*) ≥… ≥ *σ*_*n*_(*x*_0_, *t*) ≥ 0, and *U* (*x*_0_, *t*) as well as *V* (*x*_0_, *t*) are orthogonal matrices, composed column-wise of the left and right singular vectors respectively. Then

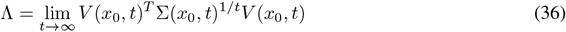

Thus, for large *t*, the log-eigenvalues of Λ can be approximated by 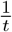 ln *σ*_*i*_(*x*_*o*_, *t*)’s, which can be thought of as the average singular value along an infinite time horizon. <u>It turns out that for ergodic systems, the Lyapunov exponents are independent of initial conditions *x*_0_. Thus, intuitively, Lyapunov exponents are topological quantities intrinsic to the dynamical system that describe the average amount of instability along infinite time horizons</u>.

In order to understand how this instability manifests along each direction, let us further look what we can say about the vectors associated with the individual Lyapunov exponents. If we denote *λ*^(1)^ ≥ *λ*^(2)^ ≥… ≥*λ*^(*s*)^ the *distinct* Lyapunov exponents, and *v*_*i*_(*x*_0_) the corresponding vector of the matrix lim_*t*→∞_ *V* (*x*_0_, *t*), then let us define the nested subspaces

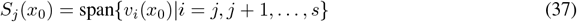

for all *j* = 1, 2, …, *s*, and take a vector *w*_*j*_(*x*_0_) ∈ *S*_*j*_(*x*_0_) \ *S*_*j*+1_(*x*_0_). Then *w*_*j*_(*x*_0_) is orthogonal to all *v*_*i*_(*x*_0_) with *i < j*, and has a non-zero projection onto *v*_*j*_(*x*_0_) since *v*_*j*_(*x*_0_) ∉ *S*_*j*+1_(*x*_0_), and thus

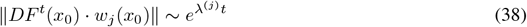

In particular, since *S*_1_(*x*_0_) is the whole phase space *X*, and *S*_2_(*x*_0_) is only a hyperplane in *X* (a subset of Lebesgue measure zero), we have that for “almost all” *w* ∈ *X*:

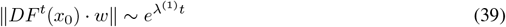

hence aligning with the direction of maximum Lyapunov exponent (MLE). In other words a randomly chosen vector, has a non-zero projection in the direction of the MLE with probability 1, and thus over time the effect of the other exponents will become negligible. This motivates taking the MLE as a way of measuring the overall amount of stability or instability of a dynamical system. One typically distinguishes the cases, where the MLE is negative, zero and positive.

Thus computing MLEs, LEs and their corresponding subspaces can be a useful tool to understand the average expansion/ contraction rate as well as the corresponding directions of gradients in recurrent neural networks.

### D.3 The QR algorithm

It is generally not advised to calculate the Lyapunov exponents and the associated vectors using *DF*^*t*^(*x*_0_) as this matrix becomes increasingly ill-conditioned. There is a known algorithm that in most cases allows to provide good estimates, called the *QR algorithm*.

As a preliminary remark, let us emphasize that the right singular vectors of *DF* (*x*_*t*+1_) do not necessarily match the left singular vectors of *DF* (*x*_*t*_), thus simply applying the singular value decomposition in order to calculate the Lyapunov exponents does not work.

Let us denote *J*_*t*_ = *DF* (*x*_*t*_) for each time step *t* = 0, 1, 2, …, then lets us pick an orthogonal matrix *Q*_0_, and compute *Z*_0_ = *J*_0_*Q*_0_. Then let us perform the QR decomposition *Z*_0_ = *Q*_1_*R*_1_. Let us further assume that *J*_0_ is invertible and we are imposing that the diagonal elements of *R*_1_ are non-negative (which we can), thus making the QR decomposition unique.

In the next step, we compute *Z*_1_ = *J*_1_*Q*_1_ and perform the QR decomposition *Z*_1_ = *Q*_2_*R*_2_, where again we are imposing the diagonal elements of *R*_2_ to be non-negative.

Continuing in this fashion at each time step *k*, we then have the identity 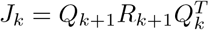, and thus

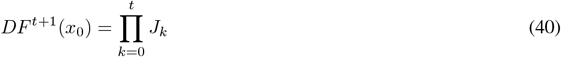

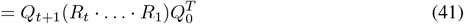

It turns out that, as long as the dynamical system is “regular”, we can then compute the *i*-th Lyapunov exponent via

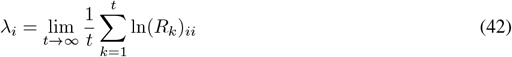

where the Lyapunov exponents are ordered *λ*_1_ ≥ *λ*_2_ ≥ … ≥ *λ*_*n*_ as explained in Benettin et al. (1980) and Dieci and Vleck (1995).

### D.4 Link to RNNs

Recalling the update equation of an RNN:

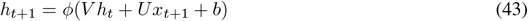

for *t* = 0, 1, …, and by denoting *F* (*h, x*) = *ϕ* (*V h* + *Ux* + *b*), we can see that

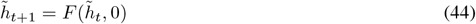

defines an autonomous discrete dynamical system (DS1), while

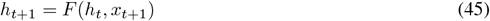

defines a non-autonomous discrete dynamical system (DS2).

For (DS1), the machinery that we have developed over the last subsections is directly applicable, as we are in the autonomous case. For instance, we can compute the Lyapunov exponents of recurrent neural network over the course of training using the QR algorithm, and in particular observe the evolution of the maximum Lyapunov exponent (MLE), as a means to measure the amount of instability or chaos in the network. For example in the case of a linear RNN with a unitary or orthogonal connectivity matrix, all LEs are equal to zero, and thus no expansion nor contraction is happening. If all LEs are negative, we are in the contracting regime, where every point eventually will approach an attractor, thus producing a vanishing gradient. For instance, Bengio et al. (1994) showed that storing information in a fixed-size state vector (as is the case in a vanilla RNN) over sufficiently long time horizon in a stable way necessarily leads to vanishing gradients when back-propagating through time (here stable means insensitive to small input perturbations).

The natural question arises whether and to what extent the machinery will stay valid for (DS2). It turns out that one can use the theory of Random Dynamical Systems Theory, where Oseledet’s multiplicative ergodic theorem holds under some stationarity assumption of the underlying distribution generating the inputs *x*_*t*_ as stated in Arnold (1998). However in this paper we are just making use of the machinery developed for (DS1), by computing Lyapunov exponents for trained RNNs but computed without inputs (*x*_*t*_ = 0 for all *t*).

## Footnotes

1 We often shorten these to *saturation* and *gain* and collectively refer to them as the **activation parameters**

